# Cellulase catalysis on cell surfaces using Caulobacter S-layer display

**DOI:** 10.1101/2025.07.28.667093

**Authors:** Beth Davenport, Tamar Av-Shalom, Kateryna Ievdokymenko, Steven J. Hallam

**Author notes:** To whom correspondence should be addressed: Dr. Steven Hallam University of British Columbia, Department of Microbiology and Immunology 2552-2350 Health Sciences Mall, Vancouver, BC Canada V6T 1Z3.

## Abstract

Sustainable bioprocesses for energy and materials production and waste resource recovery are needed to support circular economic development. Enzyme surface display on cells or functionalized materials has emerged as a promising paradigm for sustainable bioprocess innovation. Surface (S)-layers are geometric, monomolecular, highly stable crystalline protein lattices encasing the outside of many bacteria and archaea. Several S-layer genes have been shown to tolerate heterologous insertions, thereby enabling high-density display of peptides of interest on cell surfaces without additional costly immobilisation or conjugation steps. Here, we employ an S-layer display platform in *Caulobacter vibrioides* CB2A JS4038 to express functional cellulases up to 445 amino acids in length. We explore critical design considerations needed for successful catalytic display and demonstrate synergistic activities between differentially expressed cellulases relevant to combinatorial lignocellulosic biomass conversion. Functionalised S-layers capable of transforming lignocellulosic biomass could have useful applications in engineering whole-cell biocatalysts and synthetic microbial consortia tuned for different bioprocess applications.

## Introduction

Human reliance on linear resource extraction and consumption practices has resulted in cumulative negative impacts on planetary-scale operating conditions^1,2^. Increasing awareness of these impacts has led to more conscious and coordinated efforts to develop sustainable modes of production coupled to waste resource recovery laying the foundation for a circular economic model more in sync with the natural world^3^. Lignocellulosic polymers, generated through renewable photosynthetic processes, are the most abundant carbon source on earth and a promising feedstock for circular economy development^4^. For example, sixty billion tons of lignocellulosic biomass is generated annually worldwide^5^, in the form of forestry, food, crop, and plant waste, which is either transferred to landfills or burned^6^. This waste stream in isolation holds potential for generating a diverse portfolio of bioproducts^6,7^ and thus represents a frontline opportunity to circularize resource extraction with waste resource recovery^8^. Microbial communities in natural environments drive biomass conversion using lignocellulose degrading enzymes. Enzymes are widely regarded as “green”, cost-effective, highly specific catalysts that can be employed for waste resource recovery and valorisation processes^9^. For example, different types of cellulolytic enzymes, or ‘cellulases’, can be used to hydrolyse cellulose to monosaccharides for subsequent conversion into bioproducts (**Figure 1A**)^6,10^. However, many enzymes including cellulases require surface immobilisation for increased stability and catalytic efficiency to compete effectively with more energy or chemically intensive methods^8^.

**Figure 1.**
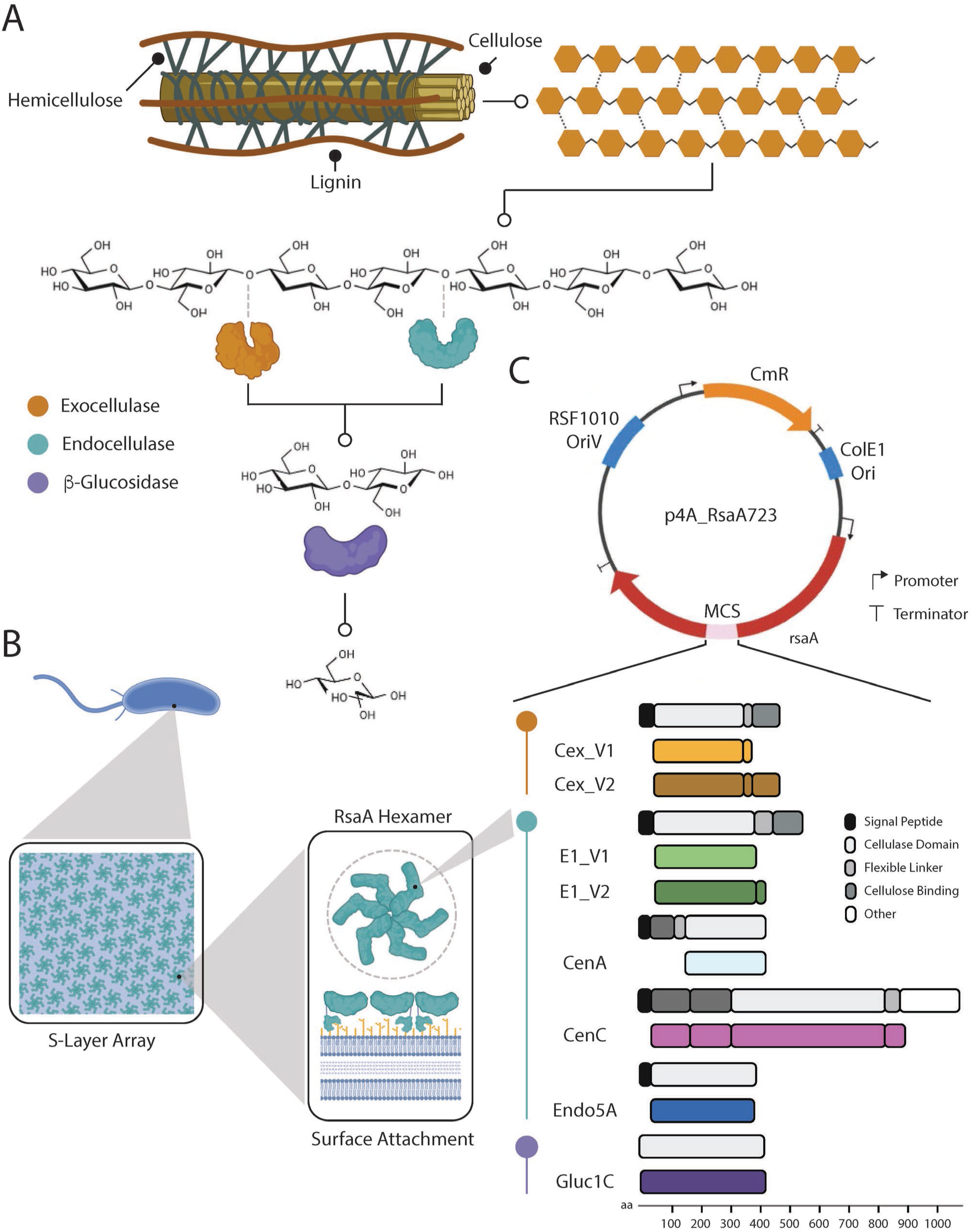
Graphical abstract illustrating the engineered S-layer display platform in Caulobacter vibrioides CB2A JS4038. A plasmid-encoded S-layer protein (RsaA) allows surface display of various cellulases, which synergistically degrade cellulose into glucose to support host growth.

Surface display has emerged as a promising biotechnology for enzyme immobilisation^9^ and prior studies using enzymes displayed on the surfaces of M13 phage, *Escherichia coli*, *Saccharomyces cerevisiae*, and *Bacillus subtilis* spores highlight the potential for microbial enzymes in waste resource recovery processes^11–14^. For example, Francisco and colleagues displayed an active cellulolytic enzyme on *E. coli* over 30 years ago^11^, Belien and colleagues demonstrated biopanning for protein–protein interactions by displaying two functional endoxylanases on M13 phage enhancing our understanding of enzyme inhibition mechanisms^15^, Guirim and colleagues co-displayed three enzymes on the surface of *S. cerevisiae* to facilitate complete conversion of Kraft Pulp to xylitol^12^, and Cho and colleagues displayed a laccase on *B. subtilis* spores to bioremediate textile dye effluent^14^. Typically, enzyme surface display involves the use of substrate binding or catalytic domains of interest translationally fused with extracellular membrane proteins^16^, which increases enzyme recyclability and combines enzyme production, purification, and immobilisation processes into a single step^8,16,17^. Current catalytic display platforms remain at early technology readiness levels due in part to prevailing limits on cell surface density and sequence insertion length^17,18^.

Bacterial and archaeal (prokaryotic) S-layers provide an adjunct or alternative platform to other display systems with potential to overcome these limitations. S-layers are geometric, monomolecular, highly stable crystalline protein lattices encasing the outside of many bacteria and archaea^19–21^. The *Caulobacter vibrioides* S-layer presents a particularly promising target for catalytic display within a genetically tractable non-pathogenic host chassis. The *C. vibrioides* S-layer is encoded by *rsaA*, which produces a constitutively expressed surface layer protein 1026 amino acids in length (∼98 kDa) secreted by a Type-1 secretion system^22,23^. Once outside of the cell, RsaA monomers self-assemble into a hexameric S-layer array composed of ∼40,000 subunits anchored to lipopolysaccharides (LPS) on *C. vibrioides’* outer membrane^19^ (**Figure 1B**). Studies have shown that *rsaA* can support insertions that enable peptide display without compromising S-layer formation^24^. Previous work by Smit and colleagues led to development of a plasmid-based S-layer display platform^25^ for in-frame *rsaA* insertion and expression in *Caulobacter vibrioides* CB2A JS4038 (**Figure 1C**). The platform was shown to support display of polypeptides up to 181 amino acids in length at 20,000 copies/μm^2^, with RsaA translational fusion protein expression reaching up to 31% of total cellular protein synthesis^26,27^. Other researchers have immobilised non-catalytic proteins up to 300 amino acids in length on the surface of the *C. vibrioides* S-layer using a linker adapter method based on chemical conjugation of the purified protein of interest to a displayed peptide tag inserted into RsaA^28,29^. However, this system requires multiple protein purification and S-layer conjugation steps with reduced display density^28^. Here, we leverage the *C. vibrioides* plasmid-based S-layer display platform developed by Smit and colleagues to constrain upper limits of protein insertion and functional display in the context of lignocellulosic biomass conversion using several different glycoside hydrolase (GH) family members including exocellulase, endocellulase, and β-glucosidase representatives (**Figure 1A-C**).

## Methods

This study encompasses three primary research activities related to media optimization, molecular cloning of surface display fusion proteins, and phenotypic characterization of expressed fusion proteins alone and in different combinations. An integrated workflow diagram is used to depict the various experimental and computational steps associated with each activity (**Figure S1**).

### Strain Selection and Growth Media Optimization

Strain sources along with relevant genotypic information are listed in **Table S1**. This study uses the S-layer display host chassis *C. vibrioides,* a heterotypic synonym of *C. crescentus,* strain JS4038 (GenBank Accession No. GCA_052446975.1), a derivative of CB2A. *C. vibrioides* CB2A JS4038 was grown in an optimised version of Peptone Yeast Extract (PYE) medium (3.2 g/L peptone, 1.8 g/L yeast extract, 0.3 g/L MgSO_4_.7H_2_0, 0.2 g/L CaCl_2_.2H_2_0) at 30°C on a culture wheel (50 rpm). Chemically competent *Escherichia coli* DH5α (Invitrogen, 18265017) was used in all intermediate cloning steps. *E. coli* was grown in Lysogeny Broth (LB) medium (1% tryptone, 0.5% yeast extract, 1% NaCl) at 37 °C with shaking at 250 rpm. When using solid medium, 1.2% (wt/vol) agar was added. 20 μg/ml chloramphenicol or 2 μg/mL chloramphenicol were used for antibiotic selection in liquid or solid media formats for JS4038 and DH5α, respectively.

PYE medium optimization was conducted using a response surface model (RSM) approach which utilised JMP Pro 18.0.1^30^, a statistical software for predictive modeling and design of experiments. The experiment involved a screen investigating the relationship between individual components of PYE medium (factors) and their impact on growth (response variable)^31–33^. To maximize growth yield, a central composite design with 5 continuous factors (Peptone, Yeast extract, CaCl_2_.2H_2_0, MgSO_4_.7H_2_0 and NaCl) was selected allowing for effective exploration of growth parameter space across 27 different media conditions (see RSM_media_components.csv), each with differing factor concentrations. Condition boundaries were set between 0 and 2 times the concentrations of each factor in previously used

PYE recipes in literature, to ensure realistic ranges conducive to *C. vibrioides* growth. 200 µL triplicates for each condition were inoculated with a starter culture of JS4038 in a round-bottom 96-well plate (Greiner, 655801) to an optical density 600 nm (OD_600_) of 0.05 and incubated at 30 °C. Growth was measured by collecting absorbance readings at 600 nm (A_600_) on a Pherastar FSX (BMG Labtech) plate reader every hour for 48 hours (**Figure S2A**). The shift between OD_600_ and A_600_ reflects the use cuvettes or plates, respectively, to estimate cell growth.

Resulting data were regressed resulting in a quadratic response surface with an R^2^ value of 0.9749 and root mean square error of 0.0689 used to identify factors associated with positive and negative growth conditions. Factors contributing to highest positive growth were Yeast extract, followed by CaCl_2_ and MgSO_4_ with NaCl identified as the primary factor contributing to negative growth (**Figure S2C**). Positive interaction terms with statistical significance indicated synergistic effects between Yeast extract x CaCl_2_, and Peptone x CaCl_2_ showing their combined influence on growth exceeded the additive effects of each factor alone. An interactive profile of cross sections of the RSM (**Figure S2D**) was used to find optimal concentrations of each factor (at desirability ‘1’) for highest growth: Peptone 3.2g/L, Yeast extract 1.8g/L, CaCl_2_.2H_2_0 0.2g/L, MgSO_4_.7H_2_0 0.3g/L. This PYE recipe was used for all following experiments in this study.

For carbon source selection experiments using carboxymethyl cellulose (CMC), a minimal medium, henceforth named Hutner’s Minimal Growth (HMG), was adapted from M6HiGG medium^34^ as follows: 1.25 mM K_2_HPO_4_, 0.025% NH_4_Cl, 1% Hutner’s Mineral Base, 0.2% carbon source (either glucose, cellobiose, or CMC). Hutner’s Mineral Base contains the following: 2% nitrilotriacetic acid, 1.5% KOH, 5.5% MgSO_4_•7H_2_0, 0.7% CaCl_2_•2H_2_0, 0.002% (NH_4_)_6_MO_7_O_24_•4H_2_0, 0.02% FeSO_4_•H20. The resulting HMG media was used to evaluate the growth of JS4038 on glucose, cellobiose, and CMC, as well as individual and combinations of RsaA:Cellulase strains on CMC.

### Sequencing Plasmid p4A_RsaA723

The plasmid p4A_RsaA723^25^ was sequenced, assembled and resolved into a single circular contig using the Oxford Nanopore platform (Plasmidsaurus). The resulting circular contig was annotated using Snapgene (v.8.1.1) (www.snapgene.com) and PlasMapper 3.0^35^ software. Manual curation was completed in the Benchling environment, using additional gene feature information derived from prior studies. A sequence-validated map of the p4A_RsaA723 plasmid backbone is represented in **Figure S3** (GenBank accession number BankIt2953359 p4A_RsaA PV578771). The plasmid includes two origins of replication, a high copy ColE1 origin for replication in *E. coli*, and a broad host range RSF1010 oriV for replication in JS4038, as well as a chloramphenicol resistance gene, and *rsaA* under the control of its wildtype promoter. This *rsaA* gene was modified to include a multi-cloning site (MCS) at amino acid position 723^36^. For downstream molecular cloning applications, p4A_RsaA723 was transformed into DH5α cells and purified with the GeneJET Plasmid Miniprep Kit (Thermo Scientific, K0503) according to the manufacturer’s instructions.

### DNA Synthesis and Molecular Cloning

Plasmids, primers, and synthesized gene blocks used in constructing RsaA fusion proteins are listed in **Table S2**, **Table S3** and **Table S4**, respectively. Six different cellulase genes were optimised for *C. vibrioides* expression using the Codon Usage Database (kazusa.or.jp) and synthesised by Thermofisher (GeneArt Custom Gene Synthesis service) or IDT (Gene Synthesis service) (**Figure 1C**). Each variant was amplified with primers that introduced BglII and PstI restriction sites for directional ligation into the p4A_RsaA723 multi-cloning site (MCS) to produce in-frame *rsaA* insertions. Q5 Hot-Start High-Fidelity polymerase (NEB, M0493) was used for DNA fragment amplification to a final concentration of 0.02 U/uL, with primers at 0.5 µM, dNTPs at 200 µM, and template DNA at 0.4 ng/uL, all in 1X Q5 Reaction Buffer. PCR cycling conditions followed the manufacturer’s protocol: an initial denaturation step at 98°C for 30 seconds, followed by 30 cycles of denaturation at 98 °C for 10 seconds, annealing at the primer-specific temperature for 30 seconds, and extension at 72 °C for 30 sec/kb, with a final extension at 72 °C for 2 min. PCR products and p4A_RsaA723 plasmid were digested with BglII (NEB, R0144) and PstI (NEB, R0140) at 37°C for 1 hr in NEBuffer r3.1, and purified with a Monarch DNA gel extraction kit (NEB, T1020) according to the manufacturer’s instructions from a 1% (w/V) agarose gel in TAE treated with SYBRSafe at 10,000x dilution (Invitrogen, S33102), and run at 90 V for 90 minutes.

Purified PCR products were ligated into p4A_RsaA723 using T4 DNA ligase (NEB, M0202) and resulting ligation products were transformed into DH5α cells according to the manufacturer’s instructions to generate colony forming units on LB agar containing appropriate antibiotics, as described above. Selected colonies were picked and grown overnight in 5 ml LB as described above. Plasmid DNA was purified with the GeneJET Plasmid Miniprep Kit (Thermo Scientific, K0503) and analysed using a Qubit 2.0 Fluorometer (Invitrogen, Q32866). Successfully assembled plasmids were confirmed using Sanger sequencing (Genewiz, Azenta Life Sciences) with primers spanning the MCS (**Table S3**) prior to transformation into JS4038. Electrocompetent JS4038 cells were prepared as described in Smit et al.^37^ In brief, 100-1000 ng plasmid DNA was transferred to 50 μl electrocompetent cells and incubated on ice for 1 min. Cells were then electroporated in a 0.1 cm gap cuvette (Bio-Rad, 1652089) with an exponential pulse of 2.5 kV, 200 Ω, 25 μF, time constant 4.2 ms using a GenePulser Xcell system (Biorad) ^37^. Cells were resuspended in 950 μl prewarmed 30°C PYE medium and incubated with shaking at 30 °C for 3 hours. Successfully electroporated cells were selected as colony forming units on PYE agar containing appropriate antibiotics, as described above.

### SDS Page and Western Blotting

Liquid 5 mL cultures for each RsaA:Cellulase expressing strain along with JS4038 and RsaA control strains were cultured for 48 hours at 30°C on a culture wheel at 50 rpm, pelleted by centrifugation for 3 min at 8,000 g and washed with 10 mM 4-(2-hydroxyethyl)-1-piperazineethanesulfonic acid (HEPES) buffer pH 7.2 prior to resuspension and dilution to OD_600_ 0.1 in HEPES. A total of 5 µL was mixed with 5 µL 2X Laemmli buffer (BioRad 1610737) containing 0.05% 2-mercaptoethanol, and then incubated at 40 °C for 10 minutes^38^. Samples were then loaded onto a BioRad 10% Stain-Free™ Protein Gel (BioRad 4568036), sandwiched between 4 µL lanes of BioRad Precision Plus Protein Standards (Bio-Rad 1610363). The gel was run at 300V for 20 minutes in SDS PAGE buffer (25 mM Tris base, 192 mM glycine, 0.1% SDS), and UV-activated for 90 seconds before imaging on a Chemidoc MP Imaging system v3.0.1.14 (Biorad). A Bio-Rad Trans-Blot Turbo system was used to transfer protein from the SDS-PAGE gel to a nitrocellulose membrane and incubated in BioRad Everyday Blocking buffer for 10 min. RsaA proteins were labeled during a 1 hour incubation with Rabbit-C Terminal Anti-RsaA polyclonal antibody^39^ (courtesy of the Smit lab) diluted 1:40,000 in Tris-Buffered Saline with 0.1% Tween-20 (TBST), followed by another 1 hour incubation with Goat-Anti Rabbit IgG StarBright™ Blue 700 diluted 1:4,000 in TBST (BioRad, 12004161). Five TBST washes were performed for 5 minutes between each incubation step. The membrane was imaged in Starbright B700 and Stain-free channels using a Chemidoc MP Imaging system v3.0.1.14 (Biorad). Successful RsaA:Cellulase protein expression was confirmed by band shift compared to the RsaA control; the relative molecular weight of bands was quantified against the BioRad Precision Plus Protein Standards, and images were brightness adjusted in ImageLab v6.1 (Biorad).

RsaA:Cellulase fusion protein surface attachment was confirmed by Western blot of low pH extracts as previously described^40,41^. In brief, 5 mL cultures for each RsaA:Cellulase expressing strain along with JS4038 and RsaA control strains were cultured for 48 hours at 30°C on a culture wheel at 50 rpm before dilution to OD_600_ 0.1 in PYE. Cells were then pelleted by centrifugation at 8,000 g for 3 minutes and washed three times with 10 mM HEPES at pH 7.2 prior to resuspension in 100 mM HEPES pH 2.0 and incubation for 10 minutes at room temperature. Samples were then neutralized with addition of 10 N NaOH followed by centrifugation at 12,000 g for 5 minutes to separate cell pellets from extracted S-layer proteins. Supernatant S-layer proteins were collected and quantified using a Pierce BCA Protein Assay Kit (Thermofisher, 23250) before dilution to 0.02-0.15 mg/mL final concentration in 10 mM HEPES pH 7.2. SDS Page and Western blotting was then performed as described above for each sample.

### Metabolic Burden and Cell Clumping Determination

Electrocompetent JS4038 cells were transformed by electroporation and selected overnight on PYE agar containing appropriate antibiotics. For metabolic burden analysis, individual colonies were transferred to 5 ml PYE containing appropriate antibiotics and cultured for 48 hours to OD_600_ 0.4–0.6 at 30 °C on a culture wheel (50 rpm). Cultures were diluted to OD_600_ 0.01 in 12 mL PYE containing appropriate antibiotics and cultured at 30 °C on a culture wheel, and OD_600_ readings were taken at periodic intervals over 30 hours. Cell clumping analysis was conducted on end time point samples based on (i) particle size distribution measurement with a MasterSizer 3000 (Malvern Panalytical) after sample sonication (50% for 30 seconds), and (ii) phase contrast light microscopy (Zeiss, standard microscope 14) at 40,000x magnification. Growth assays were repeated as above with addition of Tween-20 to the PYE media at 0%, 0.1%, 0.2%. 0.5% (Sigma-Aldrich, P1379) to reduce the cell clumping phenotype.

### Transmission Electron Microscopy

A 1% ammonium molybdate solution^42^ was prepared and pH was adjusted to 7.5 with NaOH addition. Formvar/Carbon 400 mesh Nickel grids (TED PELLA, 01754N) were made hydrophilic through glow discharge treatment. Plate colonies of JS4038 cells were gently resuspended in 10 mM HEPES Buffer pH 7.2. A nickel grid was applied to a droplet of this resuspension, incubated for 5 minutes, and excess fluid was removed with filter paper. The grid was then applied to a droplet of ammonium molybdate for 5 seconds before removing all fluid with filter paper. Transmission electron microscopy (TEM) imaging was done using a Tecnai Spirit instrument operating at 80 kV. Images were acquired at 49,000x – 68,000x using a DVC1500M side-mounted camera controlled by AMT software. This resulted in calculated pixel sizes between 1.4 nm (68,000x) – 1.9 nm (49,000x). Images were adjusted with ImageJ software^43^.

### Spectrofluorometric Cellulase Assay

A modified cellulase assay based on Mewis et al.^44^ was used to evaluate RsaA:Cellulase fusion protein function based on the production of a dinitrophenyl (DNP) chromophore released from the cellulase substrate 2,4-dinitrophenyl β-cellobioside (DNP-C) after β-1,4-glycosidic bond cleavage. Triplicate 5 mL cultures for each RsaA:Cellulase expressing strain were cultured for 48-72 hours to OD_600_ 1.0 at 30°C on a culture wheel at 50 rpm before diluting to OD_600_ 0.2 with fresh media, mixed 50:50 with assay mix to 200 µL (0.1 mg/mL DNP-C in 50 mM potassium acetate buffer at pH 5.5), loaded into a round-bottom 96-well plate (Greiner, 655801) and incubated at 30 °C. DNP-C release was measured at A_400_ absorbance using a Pherastar FSX (BMG Labtech) plate reader every hour for 24 hours, and corrected for cell growth based on concurrent A_600_ absorbance readings. The experiment was repeated using assay mixture buffer adjusted to a pH of 5, 6, and 6.5 but all functional cellulases showed highest activity at pH 5.5 as expected from prior studies with these enzymes^45–49^ (**Figure S4**).

To differentiate between cellulase activity associated with cell surfaces versus shed protein in the growth media, DNP-C assays were conducted on four different sample treatments including whole cells in PYE media, recovered PYE media, whole cells washed in 10 mM HEPES pH 7.2, and acid extracted S-layers in HEPES. Four sets of triplicate cultures expressing RsaA:E1_V1 and RsaA:E1_V2 along with JS4038 and RsaA control strains were grown to OD_600_ 1.0 at 30°C on a culture wheel at 50 rpm before diluting to OD_600_ 0.2 with fresh PYE media. These cultures we used to prepare the four different sample treatments. For whole cells in PYE media, an aliquot was taken directly for the DNP-C assay prior to centrifugation for 5 min at 4000g. After centrifugation the PYE media supernatant was collected and used for the DNP-C assay. The pellet was then washed in HEPES followed by centrifugation at 4000 g for 5 minutes three times before resuspension. The S-layer fraction was obtained from resuspended cells in HEPES as described above. For each treatment samples were mixed 50:50 with DNP-C assay mix to 200 µL (0.1 mg/mL DNP-C in 50 mM potassium acetate buffer at pH 5.5), loaded into a round-bottom 96-well plate (Greiner, 655801) and incubated at 30 °C. Plate reader measurements were then carried out as described above.

### Protein Structure Prediction

The experimentally resolved *C. vibrioides* RsaA structure was obtained from the Protein Data Bank (PDB)^50^. Structural prediction of RsaA with a multiple cloning site inserted at amino acid 723, and RsaA:Cellulase recombinant proteins was conducted using Alphafold 2^51^ on the Tamarind Bio server^52^ using default settings including the use of a single-chain model and the generation of structural ensembles with the top-ranked predicted models. Predicted structures were converted to .pdb files, which were subsequently viewed and edited in PyMOL^53^. In PyMOL, the structures were analyzed for conformational integrity, confidence scoring and visualization, employing the default rendering settings for clarity in presentation.

### Growth on Carboxymethyl Cellulose

Triplicate 5 mL cultures for RsaA:Cellulase expressing strains were cultured for 48 hours to OD_600_ 0.4–0.6 at 30 °C on a culture wheel at 50 rpm. Cultures were harvested by centrifugation at 4,500 g for 10 minutes at 4 °C, and the resulting pellets were washed in HMG three times with centrifugation at 8,000 g for 3 minutes before resuspension and dilution to OD_600_ 0.01 with HMG-carboxymethyl cellulose (CMC) media and appropriate antibiotic as described above. A total of 200 μl of diluted cells were added to a 96-well plate (Greiner, 655801) for growth and reporter assays using a Pherastar FSX plate reader (BMG Labtech) at 30 °C. To reduce evaporation, media blanks surrounded the sample wells. Absorbance at 600 nm was measured every 8 hours for seven days with orbital shaking for 5 seconds at 300rpm before readings. For synergy experiments involving two RsaA:Cellulase strains, 100 µL of each strain diluted at OD_600_ 0.01 with CMC media was added into plate wells to a final volume of 200 µL. For those involving three RsaA:Cellulase strains, 66.6 µL of each strain diluted at OD_600_ 0.01 with CMC media was added to plate wells to a final volume of 200 µL.

## Results

Caulobacter surface layer display can be realized using the p4A_RsaA723 plasmid containing *rsaA* derived from *C. vibrioides* CB15 hosted within the CB2A JS4038 Δ*rsaA* background^25^. Previous work has shown that *rsaA* tolerates in-frame DNA insertions of up to 543 nucleotides (181 amino acids) at position 2,172 (amino acid position 723), which corresponds to a surface exposed region of the protein, enabling the insertion to protrude from the multimeric hexagonal form of RsaA within the S-layer array. An MCS at this position facilitates DNA insertions using different restriction enzymes resulting in near wild type levels of expression, secretion, and surface array formation of recombinant RsaA^54,55^. We selected several well described GHs including exocellulase (EC:3.2.1.91), endocellulase (EC 3.2.1.4), and β-glucosidase (EC 3.2.1.21) representatives, which together can completely transform cellulose into monomeric sugars^56^, for in-frame insertion into *rsaA* to identify parameters including insert length, linker usage, and domain structure with potential to impact assembly of functionalized surface layer arrays (**Figure 1A-C** and **Figure S5A-C**).

### Catalytic Cellulase Display

The bifunctional beta-1,4-xylanase/glucanase exocellulase (Cex) was selected for initial evaluation because its properties and mode of action have been defined based on biochemical and crystallographic evidence^57–60^. Moreover, functional Cex has been previously expressed and secreted in *C. vibrioides* indicating compatibility with the host chassis^19^. Two different Cex variants both containing the catalytic GH domain, Cex_V1 (344 amino acids) and Cex_V2 (438 amino acids) were inserted into the p4A_RsaA723 MCS producing in-frame RsaA fusion proteins called RsaA:Cex_V1 and RsaA:Cex_V2, respectively (**Figure 1C** and **Figure S5C**). Both variants include a flexible linker after the GH domain with the potential to decrease steric hindrance and promote surface layer array formation^57,61^. In addition to catalytic domain and linker, the Cex_V2 variant also contains a cellulose binding domain (CBD) providing an opportunity to evaluate multi-domain effects on surface layer assembly and catalytic function.

Both RsaA:Cex_V1 and RsaA:Cex_V2 were expressed and secreted at high levels in the JS4038 background based on anti-RsaA immunoblotting of whole cells (**Figure 2A and S6A**). However, growth curves indicated differential metabolic burdens imposed by RsaA:Cex_V1 and RsaA:Cex_V2, respectively. No fitness cost was observed for RsaA:Cex_V1, while cells expressing RsaA:Cex_V2 grew more slowly when compared to JS4038 and RsaA controls (**Figure 2B and S7A**). Moreover, cell clumping was observed in RsaA:Cex_V2 cultures based on particle size analysis (**Figure 2C and S8A**) and confirmed using phase contrast microscopy (**Figure 2D and S8B**). The observed clumping behaviour is consistent with limited surface attachment and/or protein misfolding and aggregation^24^. Surface attachment of RsaA multimers to the cell membrane is mediated by interactions with lipopolysaccharides (LPS)^36,50^ making it is necessary to demonstrate that expressed RsaA fusion proteins are both tethered to the cell surface within intact surface layer arrays and functional with respect to known catalytic activity. Surface attachment of both RsaA:Cex_V1 and RsaA:Cex_V2 cells was evaluated using negative stain EM to determine the presence of intact surface layer arrays. Cells expressing RsaA:Cex_V1 formed intact S-layer arrays while cells expressing RsaA:Cex_V2 did not when compared to controls (**Figure 2E**). Consistent with this observation, an anti-RsaA western blot of the surface attached protein fraction indicated that RsaA:Cex_V1 was associated with the S-layer while RsaA:Cex_V2 was below the limit of detection on cell surfaces (**Figure 2F and S6B**). The impaired surface attachment of RsaA:Cex_V2 may result from steric interference between the bulky Cex_V2 domains and the N-terminal surface-anchoring region of RsaA, as suggested by AlphaFold structural predictions in comparison to RsaA:Cex_V1 or RsaA alone (**Figure 2G and S5C**). Catalytic function was determined using the DNP-C conversion assay. Interestingly, both RsaA:Cex_V1 and RsaA:Cex_V2 demonstrated cellulase activity although RsaA:Cex_V1 showed increased activity relative to RsaA:Cex_V2 (**Figure 2H and S9A**).

**Figure 2.**
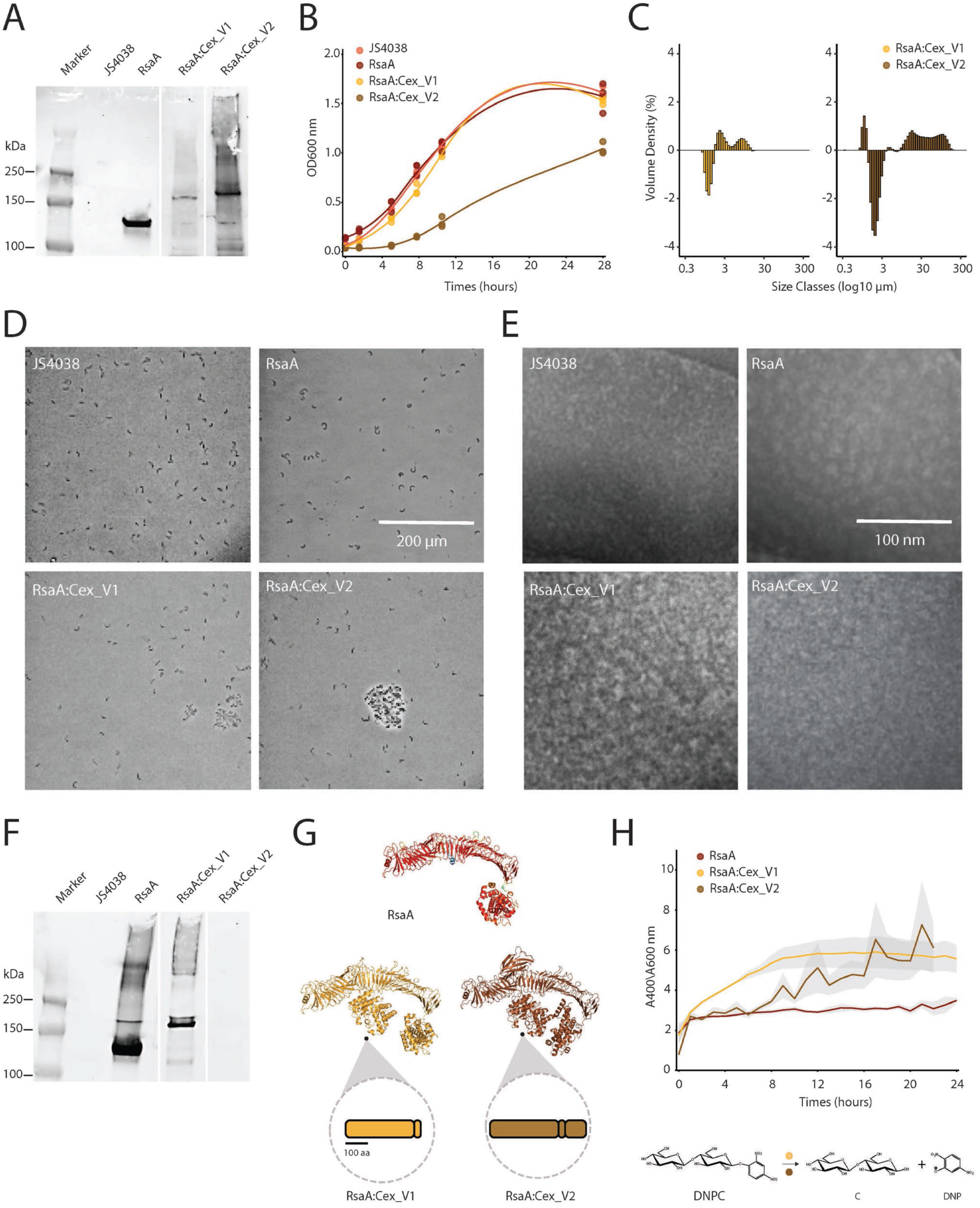
(A) Immunoblot with polyclonal anti-RsaA antibodies against whole-cells of JS4038. All lanes are from the same membrane and image. (B) Growth curves (OD_600_) of JS4038 with and without expressing RsaA or an RsaA:Cex fusion protein. Points represent individual replicates; lines represent a LOESS-smoothed best-fit line. (C) Average % volume density of four replicates of cell clump size in each strain compared to cells expressing RsaA only (% volume density of fusion strain minus % volume density of positive RsaA control at all uM), measured with a MasterSizer instrument. (D) Brightfield phase contrast microscopy images of JS4038 strains (43,000x). (E) Transmission electron microscopy images of negatively stained JS4038 strains, imaged at 49Kx mag, 80kV. (F) Immunoblot with polyclonal anti-RsaA antibodies against surface attached protein fractions of JS4038. All lanes are from the same membrane and image. (G) Alphafold predicted structures of RsaA with the multiple cloning site at amino acid 723(top), RsaA_Cex_V1 (bottom left), and RsaA_Cex_V2 (bottom right), visualised with Pymol. (H) Cellulase assay performed on JS4038 expressing RsaA or a RsaA:Cex fusion protein with DNP-Cellobioside (DNPC – bottom). Absorbance at 400 nm (A₄₀₀) was corrected for cell density using A₆₀₀. Shaded area represents the standard error of the mean of 3 biological replicates.

Based on these results, we expanded functional display efforts to include endo-1,4-β-glucanase, known as endocellulase E1, a cellulase whose properties and mode of action have been defined based on biochemical and crystallographic evidence^47,62,63^. Two different E1 variants both containing the catalytic GH domain, E1_V1 (358 amino acids) and E1_V2 (381 amino acids) were inserted into the p4A_RsaA723 MCS producing in-frame RsaA fusion proteins called RsaA:E1_V1 and RsaA:E1_V2, respectively (**Figure 1C and S5C**). RsaA:E1_V2 included a flexible linker after the GH domain with potential to decrease steric hindrance on the cell surface^57,64^. Both RsaA:E1_V1 and RsaA:E1_V2 were expressed and secreted at high levels in the JS4038 background based on anti-RsaA immunoblotting of whole cells (**Figure S6A**). No fitness cost was observed for either variant when compared to JS4038 and RsaA controls (**Figure S7B**), nor was any cell clumping evident based on particle size analysis (**Figure S8A**) or phase contrast microscopy (**Figure S8B**). An anti-RsaA Western blot of the surface attached protein fraction indicated that both RsaA:E1_V1 and RsaA:E1_V2 were associated with the S-layer albeit with RsaA:E1_V1 exhibiting much higher surface expression (**Figure S6A**). Both RsaA:E1_V1 and RsaA:E1_V2 demonstrated cellulase activity with RsaA:E1_V1 exhibiting increased activity relative to RsaA:E1_V2 (**Figure S9B**).

To further explore the relationship between insertion sequence length and formation of functional arrays, three additional endocellulases, CenA^65^, Endo5A^66^, and CenC^49^ and a β-glucosidase Gluc1C^48^ were selected for RsaA insertion without linkers. The catalytic GH domain of each protein was inserted into the p4A_RsaA723 MCS producing in-frame RsaA fusion proteins called RsaA:CenA (280 amino acids), RsaA:Endo5A (351 amino acids), RsaA:CenC (855 amino acids), and RsaA:Gluc1C (445 amino acids), respectively (**Figure 1C and S5C**). In the case of CenC, two additional cellulose binding domains were included in the fusion protein. Both RsaA:CenA and RsaA:Endo5A were expressed and secreted based on anti-RsaA immunoblotting of whole cells (**Figure S6A**). No fitness cost was observed for either fusion protein when compared to JS4038 and RsaA controls (**Figure S7B**), nor was any cell clumping evident based on particle size analysis (**Figure S8A**) or phase contrast microscopy (**Figure S8B**). An anti-RsaA Western blot of the surface attached protein fraction indicated that neither RsaA:CenA and RsaA:Endo5A were associated with the S-layer similar to results observed for exocellulase RsaA:Cex_V2 (**Figure S6B**). Both fusion proteins exhibited cellulase activity using the DNP-C conversion assay (**Figure S9C**). The multi-domain insert RsaA:CenC was expressed and secreted at high levels based on anti-RsaA immunoblotting of whole cells (**Figure S6A**). However, the observed protein appeared truncated at 95 kDa indicating possible internal cleavage or cryptic translation initiation artefacts. Moreover, RsaA:CenC imposed a metabolic burden when compared to JS4038 and RsaA controls (**Figure S7C**) and exhibited extreme cell clumping in culture (**Figure S7A-B**). Similar to RsaA:CenA and RsaA:Endo5A an anti-RsaA Western blot of the surface attached protein fraction indicated that RsaA:CenC was not associated with the cell surface (**Figure S6B**). Moreover, the fusion protein did not exhibit cellulase activity using the DNP-C conversion assay (**Figure S9D**). RsaA:Gluc1C was expressed at low levels by JS4038 cells (**Figure S6A**), imposed minimal metabolic burden (**Figure S7D**) with no clumping phenotype (**Figure S8A-B**), and exhibited some surface attachment (**Figure S6B**). Despite limited expression within cells and on cell surfaces RsaA:Gluc1C retained cellulase activity using the DNP-C conversion assay (**Figure S9D**) making it the longest fusion protein successfully displayed on the surface of *C. vibrioides* cells.

Current conception of the S-layer life cycle considers RsaA monomers as continuously transcribed, translated, secreted and added to existing S-layer crystals, especially at points of localized defects or extreme topology e.g., cell poles, division planes and stalk or flagella bases^67^. This dynamic model indicates a potential for monomer turnover and shedding into the surrounding media which in turn could confound interpretation of functional assays for surface activity. To differentiate between the activity of functional surfaces and soluble media components during the S-layer life cycle RsaA:E1_V1 and RsaA:E1_V2 activity was measured on 1) unwashed cells in PYE media which should include all displayed S-layer as well as any shed S-layer during cell growth and replication, 2) washed cells in HEPES buffer, removing all previously shed S-layer present in the media, 3) surface attached protein fraction only, obtained using low pH extraction, and 4) PYE media without cells from treatment 1) containing shed S-layer components (**Figure 3A**).

**Figure 3.**
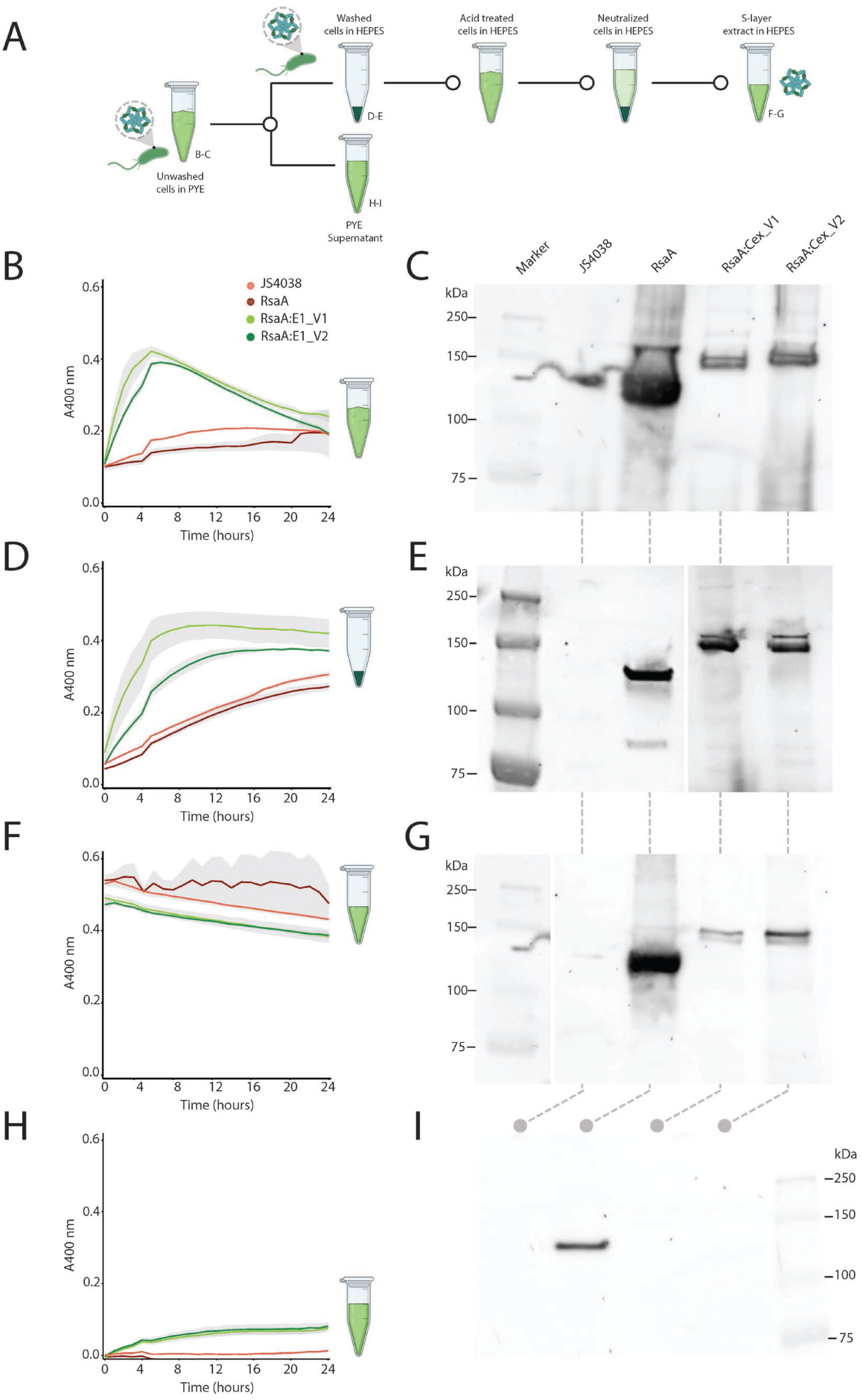
Comparison of cellulase activity from surface attached protein vs. that shed into the media. (A) Experimental design workflow (B, D, F, H) DNP-C cellulase assays with error bars representing standard error of the mean of three biological replicates for treatment 1) (unwashed JS4038 in PYE), 2) (washed JS4038 in HEPES buffer), 3) (surface attached protein fractions), and 4) (PYE culture media only from treatment 1) respectively. (C, E, G, I) Accompanying immunoblots with polyclonal anti-RsaA antibodies of treatment 1), 2), 3) and 4) respectively. Immunoblot from treatment 2) performed on a separate gel to those from treatment 1), 3) and 4).

Cellulase activity for RsaA:E1_V1 and RsaA:E1_V2 in treatment 1) reached peak A400 absorbance values of 0.421 and 0.390 (**Figure 3B**), respectively while cellulase activity for RsaA:E1_V1 and RsaA:E1_V2 in treatment 2) reached peak A400 absorbance values of 0.442 and 0.377, respectively consistent with functionalized cell surfaces (**Figure 3D**). No cellulase activity was measurable for treatment 3) due to low pH treatment used to remove the S-layer from cell surfaces resulting in a loss of enzyme activity (**Figure 3F)**. However, cellulase activity in the media from treatment 1 reached peak A400 absorbance values of 0.0765 and 0.0815 for RsaA:E1_V1 and RsaA:E1_V2 respectively, approximately 5-fold lower than treatment 1) or 2) (**Figure 3F)**. These results were reinforced using western blotting for each treatment indicating high levels of RsaA:E1_V1 and RsaA:E1_V2 displayed on the cell surface, with little to no protein detected in PYE media (**Figure 3C, E, G and I**). Taken together these results indicate that while most of the activity associated with RsaA:E1_V1 and RsaA:E1_V2 was associated with cell surfaces, a small fraction of shed S-layer present in the media can contribute to measured activity consistent with the dynamic model.

### Practical Design Considerations

The availability of X-ray crystallographic and electron cryotomography (cryo-ET) information for *C. vibrioides* RsaA provides a baseline for predictive comparisons^50,68^. Based on 2.7 Å X-ray and 7.4 Å cryo-ET resolved structural information, six RsaA monomers assemble into a hexagonal array ∼240 Å in length coalescing around a central pore ∼20 Å in width^68^. Visualized in PyMol the structure for RsaA appears as an L-shaped monomer with several important features resolved, including an N-terminal LPS binding domain needed to anchor RsaA arrays to the cell surface, and several internal calcium (Ca^2+^) binding regions necessary for stable array formation (**Figure S5A**). The predicted RsaA723 structure with intact MCS corresponds closely to wild type RsaA providing a sanity check on Alphafold2 results (**Figure S5A-B**). Building on this information, Alphafold2 was used to predict three-dimensional structures for RsaA:Cex_V1, RsaA:Cex_V2, RsaA:E1_V1, RsaA:E1_V2, RsaA:CenA, RsaA:Endo5A, RsaA:CenC, and RsaA:Gluc1C (**Figure S5C**) and the resulting predictions were then used to assess potential impacts on observed phenotypic characteristics (**Table 1**).

**Table 1.** Summary of design features and phenotypic characteristics of surface displayed parts. Design features can be used to assess potential impacts on observed phenotypic characteristics of displayed parts.

|  | Surface Displayed Parts |  |  |  |  |  |  |  |  |
| --- | --- | --- | --- | --- | --- | --- | --- | --- | --- |
|  | RsaA | CenA | CenC | Cex_V1 | Cex_V2 | E1_V1 | E1_V2 | Endo5A | Gluc1C |
| **Design Feature** |  |  |  |  |  |  |  |  |  |
| Protein length (aa) | 1026 | 280 | 855 | 340 | 438 | 358 | 381 | 352 | 445 |
| Molecular weight (kDa) | 98 | 30 | 90 | 37 | 47 | 40 | 42 | 44 | 51 |
| Isoelectric point (pI) | 3.8 | 6.4 | 4.1 | 5 | 5.1 | 4.9 | 4.9 | 5.3 | 5 |
| Intramolecular disulfide bonds | 0 | 2 | 4 | 2 | 3 | 2 | 2 | 0 | 1 |
| Domain number | 2 | 1 | 4 | 1 | 2 | 1 | 1 | 1 | 1 |
| Structurally predicted steric hindrance | 0 | 0 | 1 | 0 | 1 | 0 | 0 | 0 | 0 |
| **Phenotype** |  |  |  |  |  |  |  |  |  |
| High level expression | 1 | 1 | 0 | 1 | 1 | 1 | 1 | 1 | 0 |
| Cellulase activity | 0 | 1 | 0 | 1 | 1 | 1 | 1 | 1 | 1 |
| Surface attachment | 1 | 0 | 0 | 1 | 0 | 1 | 1 | 0 | 1 |
| Metabolic burden | 0 | 0 | 1 | 0 | 1 | 0 | 0 | 0 | 0 |
| Cell clumping | 0 | 0 | 1 | 0 | 1 | 0 | 0 | 0 | 0 |

While RsaA:Cex_V1 was predicted to fold correctly with minimal attachment interference consistent with observed array formation and cellulolytic activity, RsaA:Cex_V2 appeared to introduce steric hindrance to the N-terminal RsaA domain essential for LPS attachment. This hindrance likely arises from the increased spatial demands required for the proper projection of both the Cex_V2 CBD and catalytic domain (**Figure S5C**). These predicted changes are consistent with the observed clumping and accumulation of RsaA:Cex_V2 in the growth medium. Predictions for RsaA:E1_V1 and RsaA:E1_V2 also indicated correct folding with minimal attachment interference consistent with observed array formation and activity (**Figure S5C**). Interestingly, both RsaA:CenA and RsaA:Endo5A were predicted to fold correctly despite exhibiting no array formation based on anti-RsaA Western blot of the surface attached protein fraction (**Figure S5C**). Consistent with phenotypic results indicating a clumping and truncated product, the multi-domain insert RsaA:CenC was predicted to distort β-strands near the MCS associated with Ca^2+^ binding and interfere with RsaA N-terminal surface attachment on folding (**Figure S5C**). The predicted RsaA:Gluc1C structure indicated correct folding (**Figure S5C**), also consistent with phenotypic and Western blot data as surface attaching, representing the longest cellulase successfully displayed on this platform at 445 amino acids.

Given the combined results, insertion sequence length in isolation is not predictive of functional array formation, although increasing the length and domain complexity of RsaA fusion proteins likely imposes pleiotropic effects on gene expression, protein folding, stability, and secretion with the potential to impede cell growth and limit or inhibit functional array formation on the surface of *C. vibrioides* cells. Despite this uncertainty, the use of Alphafold2 or other structure prediction algorithms does provide a basis for comparing sequence features in relation to expected functional outcomes. Notably, predicted structural steric hindrance correlates with constructs exhibiting cell clumping and metabolic burden phenotypes (**Figure S10**), which represent the most informative phenotypic markers of failed display in this study. This approach is useful in optimizing functional display and in the design of combinatorial growth experiments.

### Synergistic Effects

In nature, combinations of microbially encoded exocellulases, endocellulases, and β-glucosidases can synergistically transform cellulose into monomeric sugars^56^ to support cell growth and renewal. Such emergent properties where the whole is greater than the sum of its parts remain integral to microbial community metabolism and present an opportunity to learn from nature in the design of microbial cell factories and synthetic consortia tuned for biomass transformation applications. In this light, S-layer display could provide an organizing principle for development of emergent processes in which ensembles of cells cooperate to transform biomass into useful bioproducts. To test this idea, growth experiments were conducted using different combinations of functionally displayed cellulases including RsaA:Cex_V1, RsaA:E1_V2, and RsaA:Gluc1C described above to promote growth of JS4038 on CMC as the sole carbon source (**Figure 4A-B** and **Figure S11**).

**Figure 4.**
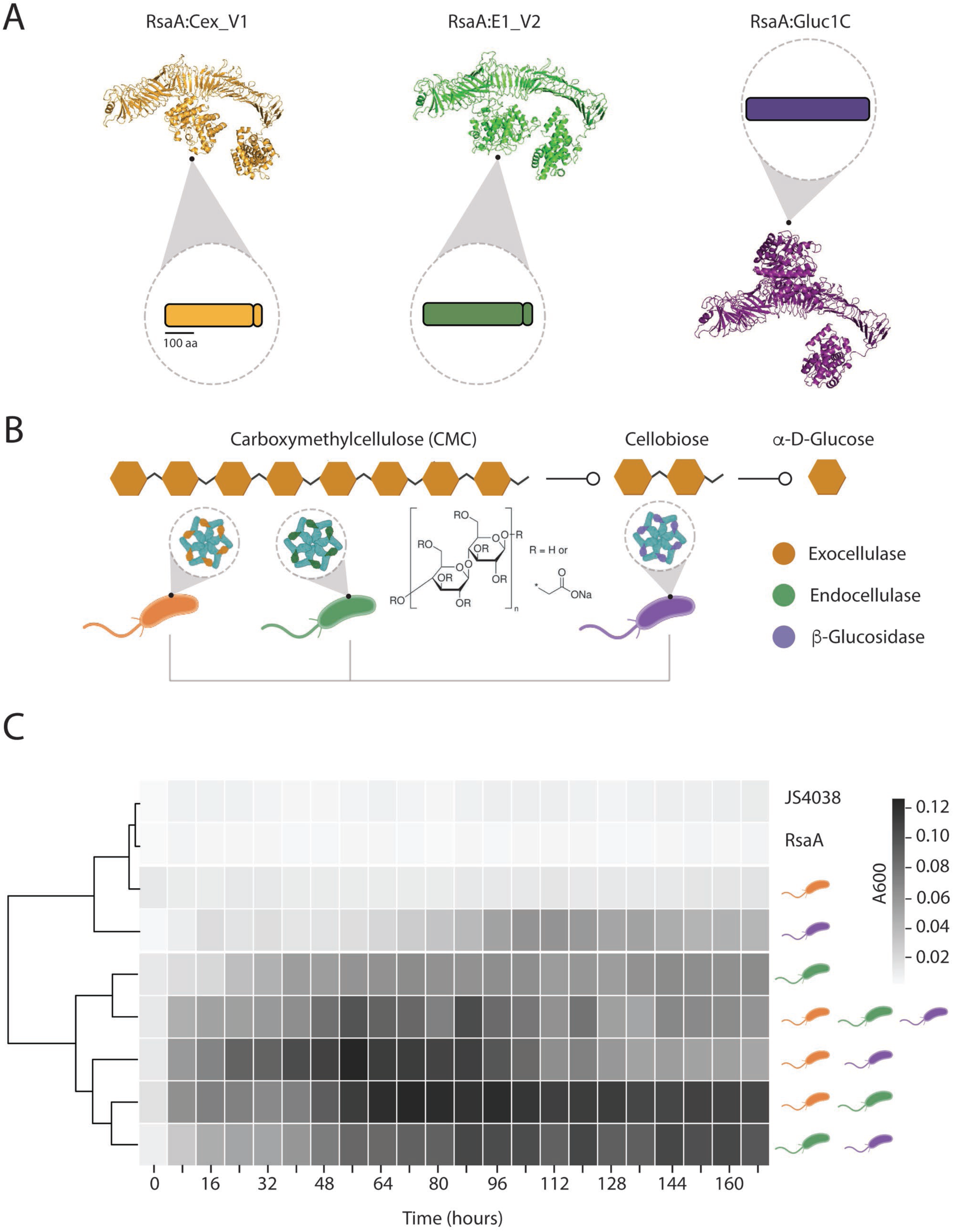
(A) Alphafold structural models of three RsaA fusion proteins used in the CMC growth study. (B) Schematic showing where the three cellulase classes cleave to break down CMC to glucose. (C) Heatmap representing mean growth (A_600_) of three biological replicates of JS4038 expressing RsaA:Cellulase fusion proteins in HMG minimal media with CMC as the sole carbon source. Rows represent individual strains or mixtures of strains, with hierarchical clustering visualised on the left.

Strain JS4038 cannot catabolize CMC presenting an opportunity for gain of function experiments. However, before testing, an alternative medium needed to be concocted supporting JS4038 growth under carbon source selection. Although previous studies have reported using modified M2 minimal media for Caulobacter growth^69,70^, initial experiments with M2 were unsuccessful prompting development of an alternative minimal medium called HMG (see methods). JS4038 grew to densities exceeding OD_600_ 0.5 in HMG supplemented with glucose, to cell densities approaching OD_600_ 0.15 on cellobiose (JS4038 has some endogenous β-glucosidase activity^71^), and not at all on CMC (Figure 4C **Figure S11B**). However, JS4038 expressing either RsaA:Cex_V1, RsaA:E1_V2 or RsaA:Gluc1C marginally grew on CMC approaching OD_600_ 0.01, 0.06, and 0.06, respectively reaching saturation between 60-100 hours (**Figure 4C and S11C-E**). When different combinations of RsaA:Cex_V1, RsaA:E1_V2, or RsaA:Gluc1C were grown together, higher cell densities were realized (**Figure 4C and S11C-E**). For example, a 50:50 mix of RsaA:Cex_V1 and RsaA:Gluc1C enabled JS4038 to reach OD_600_ 0.12 saturating within 50 hours (**Figure 4C and S11C**). Similarly, a 50:50 mix of RsaA:Cex_V1 and RsaA:E1_V2 enabled JS4038 to reach OD_600_ 0.12 saturating within 70 hours (**Figure 4C and S11D**), while a a 50:50 mix of RsaA:E1_V2 and RsaA:Gluc1C enabled JS4038 to reach OD_600_ 0.10 saturating within 100 hours (**Figure 4C and S11E**). Interestingly a mixture of RsaA:Cex_V1, RsaA:E1_V2, and RsaA:Gluc1C in equal proportions enabled JS4038 to reach a maximum of OD_600_ 0.12 within 70 hrs before declining to OD_600_ 0.06 between 70-170 hours (**Figure 4C and S11E**). Consistent with their enhanced growth phenotypes, the time-resolved growth profiles of the JS4038 mixtures clustered closely under hierarchical clustering analysis (**Figure 4C**). It is interesting to note that the addition of RsaA:Gluc1C to a mixture of strains RsaA:E1_V2 and RsaA:Cex_V1 decreases the CMC growth phenotype; This effect may reflect product inhibition, as many β-glucosidases are known to be feedback-inhibited by glucose^72^. These results indicate the potential for synergistic transformation of cellulosic feedstocks using synthetic consortia expressing functionalized cell surfaces containing different GH combinations.

## Conclusion

This study demonstrates that it is possible to insert intact or nearly intact functional proteins up to 445 amino acids in length into RsaA monomers, and that resulting fusion proteins can form arrays that retain catalytic function on the surface of *C. vibrioides* cells. In all instances, linker addition did not appear to reduce steric hindrance needed for functional array formation and can as in the case of RsaA:E1_V2 reduce expression or activity of displayed proteins. Moreover, proteins containing multiple domains tended to clump together, limiting or preventing surface attachment and functional array formation. Combinatorial growth experiments indicated that different cellulase combinations displayed on cell surfaces can support growth on CMC, and that this growth can in some instances be synergistic as in the case of RsaA:Cex_V1 and RsaA:E1_V2. Unexpectedly, cell mixtures displaying exocellulase, endocellulase, and β-glucosidase in equal proportions did not necessarily improve this growth trajectory. This could be the result of multiple factors including but not limited to incomplete substrate conversion, end-product inhibition, toxic intermediate production, or differential fitness effects between strains harboring alternative surface display constructs during co-culture. It would be of further interest to explore these factors in more detail in relation to the development of whole-cell biocatalysts and microbial consortia tuned for the breakdown and funneling of different lignocellulosic feedstocks and in the design of novel screening paradigms for recovery of surface displayed catalytic functions from environmental genomes. Moreover, although the use of linkers did not appear to improve functional array formation, the concept of combinatorial attachment using differential linker display in co-culture could provide a scaffold for assembly of more complex multi-component enzyme complexes on *C. vibrioides* cell surfaces supporting an even wider range of bioprocess applications. Future experiments could explore co-culture between functionalized JS4038 surfaces and other strains tuned for lignin- or hemicellulose transformation or with strains engineered to produce value-added compounds from JS4038 funneled substrates providing a novel framework for developing synthetic microbial consortia with functionalized cell surfaces.

## Acknowledgements

We would like to thank John Smit for his practical wisdom and consistent support in working with *C. vibrioides* and the plasmid-based display system, all the members of the 2016 University of British Columbia International Genetically Engineered Machine (iGEM) team, who first conceived of the idea to display cellulases on the surface of *C. crescentus*, and John Nomellini, who helped with original construct design. Additionally, we would like to acknowledge Kit Martens, who cloned two of the plasmids used in this study. Finally, we would like to thank Nicholas Lin for providing critical feedback on the manuscript. This work was performed under the auspices of the Natural Sciences and Engineering Research Council (NSERC) of Canada, the Canada Foundation for Innovation (CFI), Genome British Columbia, the Rio Tinto Centre for Future Materials, and the BRIMM Mining Microbiome Theme, with essential automation support through the Biofactorial high-throughput biology facility in the Life Sciences Institute at the University of British Columbia. BD was supported by the Four Year Doctoral Fellowship (4YF) program at the University of British Columbia, the Dr. Linda Matsuuchi Graduate Scholarship in Life Sciences, and the Rio Tinto Graduate Scholarship.

