## Supplementary Figures and Tables for "Cellulase catalysis on cell surfaces using Caulobacter S-layer display"

**Table S1** Strain sources and genotypic information.

| Name | Description | Genotype | Source | NCBI Genome Accession no. |
|---|---|---|---|---|
| *Escherichia coli* DH5α | High efficiency cloning strain | F– ϕ80lacZΔ M15 Δ (*lacZYA-argF*) *U169 recA1 endA1 hsdR17* (rK– mK+) *phoA supE44* λ- *thi–1 gyrA96 relA1* | Thermofisher (cat: EC0112) | GCF_000982435.1 |
| *Caulobacter vibrioides CB2A JS4038* | Chassis strain adapted for the S-layer display system contains point mutation in *rsaA* locus, deletion in S-layer associated protease *sap* locus, BAC genes integrated into *xylX* locus, and deletion of GDP-L-fucose synthase *fcl* locus involved in EPS synthesis | *ΔrsaA ΔsapA Δfcl ΔxylX::repBAC* | Farr et al. (2013), Davenport et al. (2025) | GCA_052446975.1 |

**Table S2** Plasmids used in this study.

| Name | Description | Resistance Marker | Source |
|---|---|---|---|
| p4A_RsaA | CB15 *rsaA* with multiple cloning site at amino acid 723 | Chloramphenicol | Nomellini et al. (2007) |
| p4A_RsaA_CenA | CB15 *rsaA* with CenA cellulase insert | Chloramphenicol | This study |
| p4A_RsaA_CenC | CB15 *rsaA* with CenC cellulase insert | Chloramphenicol | This study |
| p4A_RsaA_Cex_V1 | CB15 *rsaA* with Cex cellulase insert | Chloramphenicol | This study |
| p4A_RsaA_Cex_V2 | CB15 *rsaA* with Cex cellulase insert | Chloramphenicol | This study |
| p4A_RsaA_E1_V1 | CB15 *rsaA* with E1 cellulase insert | | This study |

| | | | |
|---|---|---|---|
| | | Chloramphenicol | |
| p4A_RsaA_E1_V2 | CB15 *rsaA* with E1 cellulase insert | Chloramphenicol | This study |
| p4A_RsaA_Endo5A | CB15 *rsaA* with Endo5A cellulase insert | Chloramphenicol | This study |
| p4A_RsaA_Gluc1C | CB15 *rsaA* with Gluc1C cellulase insert | Chloramphenicol | This study |

**Table S3** Primers used in this study.

| **Name** | **Sequence (5' > 3')** | **Source** |
|---|---|---|
| MCS_fwd | CGACGGCACCGACGTTCT | This study |
| MCS_rev | CGGAGCCGCCAGAACGGTCAGGCCGACATTCAC | This study |
| CenA_fwd | TCCAGATCT**CAGCCGACGAGCGGGTTTTACGTTG** | This study |
| CenA_rev | TCCATCTGCAGA**CCAGCGCGCGTTACGAGCCATTTC** | This study |
| CenC_fwd | TCCAGATCT**AGCACGGGAGCATTGTGCGTTGCG** | This study |
| CenC_rev | TCCATCTGCAGA**CTCAGCGCTCCCCTGATCGGCAAC** | This study |
| Cex_V1_fwd | GGGAGATCT**GCGACCACGCTCAAGGAGGCCGCC** | This study |
| Cex_V1_rev | CTAGCTAGCC**CCGGCCGGACCGGACGTCGG** | This study |
| Cex_V2_fwd | TCCAGATCT**GCCACGACCCTGAAAGAAGC** | This study |
| Cex_V2_rev | TCCATCTGCAGA**CGTGCCGTTCAAGCTAAAAGC** | This study |

| | | |
|---|---|---|
| E1_V1_fwd | TCCAGATCT**GTTGCAGGCGGGGGTTATTG** | This study |
| E1_V1_rev | TCCATCTGCAGA**AACCGGGTCAAATATCGATGATTTTATC** | This study |
| E1_V2_fwd | TCCAGATCT**GTTGCAGGCGGGGGTTATTG** | This study |
| E1_V2_rev | TCCATCTGCAGA**TGAGGGGGAGGGAGAC** | This study |
| Endo5A_fwd | TCCAGATCT**AGCGTCAAGGGGTATTACCAC** | This study |
| Endo5A_rev | TCCATCTGCAGA**CACCGGCTTCATGATCCG** | This study |
| Gluc1C_fwd | TCCAGATCT**AACACGTTCATCTTTCCGGC** | This study |
| Gluc1C_rev | TCCATCTGCAGA**GAACCCGTTCTTGGCCAT** | This study |

**Table S4** DNA parts used in this study.

| **Name and Truncations** | **Description** | **Source** | **Sequence** |
|---|---|---|---|
| CenA<br><br>(CenA – 840nt, 280aa) | *Cellulomonas fimi;* endo- β - 1,4-glucanase, Uniprot: P07984 | Wong et al. (1986) | ATGAGCACCCGTCGCACGGCAGCAGCACTGTTGGCAGCGGCAGCTGTTGCCGTCGGAGGGCTGACCGCTCTTACGACCACGGCGGCCCAGGCGGCTCCAGGCTGCCGCGTTGATTATGCAGTCACCAATCAGTGGCCGGGTGGTTTCGGAGCAAACGTGACCATCACGAATCTGGGGGACCCCGTCTCGTCGTGGAAACTGGACTGGACCTACACCGCGGGCCAGCGCATCCAACAACTGTGGAACGGAACGGCAAGCACCAACGGTGGGCAGGTGAGCGTTACCTCGTTGCCTTGGAACGGCTCGATTCCGACCGGAGGGACGGCTTCCTTTGGCTTTAACGGTTCCTGGGCAGGTTCCAATCCAACGCCAGCGAGCTTCTCCTTGAACGGGACCACCTGCACGGGCACGGTTCCAACCACGAGCCCTACTCCTACGCCGACCCCCACCACTCCGACACCGACGCCGACTCCGACGCCTACCCCAACCCCTACGGTTACGCCCCAGCCGACGAGCGGGTTTTACGTTGATCCTACGACCCAAGGATATCGTGCTTGGCAGGCCGCAAGCGGTACGGACAAGGCCCTGTTGGAGAAGATCGCGCTTACCCCACAAGCATATTGGGTCGGGAACTGGGCTGATGCAAGCCACGCTCAAGCAGAGGTTGCGGATTACACCGGGCGCGCGGTGGCAGCCGGGAAGACGCCCATGTTGGTCGTGTACGCAATCCCGGGTCGTGACTGCGGCAGCCACAGCGGAGGGGGCGTGTCGGAGTCCGAATACGCGCGTTGGGTGGATACGGTGGCCCAAGGTATTAAGGGGAATCCGATTGTGATCCTGGAGCCTGACGCTCTGGCTCAACTGGGAGACTGCTCGGGGCAAGGCGACCGTGTGGGGTTTCTGAAATATGCCGCTAAAAGCCTTACCCTTAAAGGGGCTCGTGTCTACATTG |

| | | | |
|---|---|---|---|
| | | | ACGCCGGTCATGCCAAATGGTTGAGCGTTGATACCCCGGTTAATCGC<br>TTGAATCAGGTGGGTTTTGAGTATGCTGTGGGATTTGCCTTGAACACG<br>AGCAACTATCAAACGACCGCGGACTCCAAGGCTTATGGTCAGCAAA<br>TCTCCCAACGCCTTGGCGGTAAGAAGTTTGTGATCGATACGTCGCGC<br>AACGGCAACGGGAGCAACGGGGAGTGGTGCAATCCTCGTGGTCGT<br>GCTTTGGGAGAACGTCCCGTGGCAGTTAACGACGGAAGCGGTTTGG<br>ACGCCCTGTTGTGGGTGAAACTGCCCGGAGAGAGCGATGGCGCAT<br>GCAACGGTGGACCTGCTGCTGGTCAGTGGTGGCAAGAGATCGCGC<br>TGGAAATGGCTCGTAACGCGCGCTGG |
| CenC<br><br>(CenC – 2565nt, 855aa) | *Cellulomonas fimi;* endo-β-1,4-glucanase and exo-β-1,4-glucanase, Uniprot: P14090 | Tomme et al. (1996) | ATGGTAAGCCGCCGCAGTTCTCAAGCACGCGGTGCGTTAACCGCA<br>GTCGTTGCTACTTTAGCCCTTGCCTTAGCCGGATCAGGTACGGCCCT<br>GGCGGCGAGTCCCATCGGCGAGGGGACATTTGACGATGGTCCAGA<br>GGGATGGGTCGCATACGGTACAGATGGCCCATTAGACACAAGCACG<br>GGAGCATTGTGCGTTGCGGTACCGGCAGGATCAGCTCAATATGGAGT<br>CGGCGTCGTACTTAACGGGGTCGCTATCGAAGAGGGTACCACTTATA<br>CACTGCGTTACACCGCTACGGCCAGTACAGACGTAACAGTGCGCG<br>CGTTAGTTGGCCAGAACGGCGCTCCGTACGGGACTGTTCTTGATAC<br>GAGCCCGGCCTTGACAAGCGAACCACGTCAAGTTACGGAAACCTTC<br>ACTGCAAGTGCCACGTACCCGGCCACGCCCGCCGCAGACGATCC<br>TGAGGGTCAGATTGCTTTCCAATTGGGTGGGTTCTCCGCCGACGCAT<br>GGACTTTCTGTCTGGACGACGTCGCTTTAGACTCTGAGGTAGAGTTAT<br>TGCCTCATACATCCTTTGCTGAATCGCTGGGCCCGTGGAGTCTTTAC<br>GGGACTAGCGAGCCCGTGTTTGCCGATGGGCGCATGTGCGTAGACT<br>TGCCCGGAGGCCAAGGGAATCCCTGGGACGCAGGACTGGTATATA<br>ACGGTGTCCCGGTAGGTGAGGGGGAGAGTTATGTATTGAGCTTCACA<br>GCGTCGGCCACGCCTGACATGCCGGTACGCGTATTAGTAGGTGAAG<br>GGGGAGGGGCATATCGCACAGCGTTTGAGCAAGGGAGTGCCCCG<br>CTGACAGGCGAGCCTGCCACACGCGAATATGCTTTTACGTCTAATTT<br>GACATTTCCGCCTGATGGGGACGCACCTGGCCAGGTAGCATTTCAC<br>CTTGGGAAGGCGGGCGCCTACGAGTTTTGTATTAGTCAGGTGAGCTT<br>GACGACATCGGCTACTCCTCCCCCCGGTTATGAACCAGATACCGGC<br>CCACGTGTCCGCGTTAATCAAGTCGGCTATTTACCGTTCGGGCCGAA<br>ACGTGCCACGCTTGTAACGGACGCGGCCGAGCCCGTCGCATGGG<br>AATTGCGTGACGCAGATGGTGTGGTCGTGGCAGATGGCACATCAGAA<br>CCTCGTGGCGTAGAGCCTTCTGCCGCGCAGGCGGTCCACGTCTTG<br>GATTTCTCGGACGTCACGACTCAGGGCGCGGGATACACGTTAGTTG<br>CCGACGGAGAAACGAGCCGTCCCTTTGATATCGATGGAGACCTTTAC<br>CAGCAGTTGCGCTACGACGCGTTGAATTACTTTTATCTTGCGCGTAGT<br>GGCACAGAAATCGAGGCGGATGTCGTCGGGGAGGAGTACGCGCG<br>CGAAGCGGGCCATGTCGGAGTAGCTCCTAACCAGGGAGACACTGA<br>CGTGCCGTGCATTGGTCCCCGCGATTATTATGACGGGTGGACATGTG<br>ACTACCGTTTGGATGTATCTGGCGGCTGGTATGACGCAGGCGACCA<br>CGGAAAATACGTCGTAAACGGAGGGATTGCGGTAGGGCAATTGCTTC<br>AAACATACGAGCGCGCTCTGCATGCAGGTACCGCGGATGCGCTTG<br>CAGATGGTACTTTAGACGTACCGGAACATGGCAATGATGTGCCGGAC<br>GTGTTAGATGAAGCGCGCTGGGAATTGGAGTGGATGTTATCTATGATT<br>GTCCCTGAGGGAGAGTATGCCGGGATGGTACATCATAAAGTGCACGA<br>CGAGGGATGGACAGGTCTTCCCTTACTTCCCGCTGACGACCCACAA<br>GCCCGCAGCTTACACCGTCCTAGCACAGCGGCGACGCTGAATCTG<br>TCGGCAGTAGCTGCACAAGGCGCGCGTTTGTTGGAACCATACGATC<br>CACAATTAGCTCAGACACTGTTGGAGGCCGCGCGCACAACTTGGGC<br>AGCAGCTCAAGAGCATCCCGCGCTGTATGCCCCTGGTGAGGCCGG |

| | | | |
|---|---|---|---|
| | | | AGCTGATGGAGGTGGTGCTTATAATGATTCGCAAGTGGCAGATGAATT<br>TTACTGGGCGGCTGCCGAGCTGTATTTGACAACCGGGGAGGACGC<br>ATTTGCCACAGCCGTGACCACCTCCCCATTACATACAGCCGATGTCT<br>TCACTGCGGATGGGTTCGGTTGGGGCTCGGTTGCCGCATTAGGACG<br>CTTGGACTTGGCAACAGTCCCGAACGAGTTACCTGGTTTAGATGCTG<br>TACAGAGCTCGGTCGTCGAGGGGGCACAAGAGTACCTTGCAGCTCA<br>AGCCGGGCAGGGATTTGGATCACTTTACAGTCCCCCGGGTGGTGAA<br>TACGTGTGGGGATCGAGTTCGCAGGTCGCGAATAATCTTGTAGTGGT<br>GGCCACCGCTTACGATTTAACTGGAGATGAGCGCTTTCGTGCAGCAA<br>CGCTGGAGGGGTTGGACTATTTGTTCGGCCGTAACGCCTTAAATCAG<br>TCTTACGTTACTGGCTGGGGAGAGGTTGCATCTCATCAACAGCACTC<br>GCGCTGGTTTGCTCATCAGTTGGACCCGTCACTTCCAAGTCCGCCG<br>CCGGGTAGCTTGGCAGGAGGCCCTAATTCTCAGGCCGCTACTTGGG<br>ACCCAACGACTAAAGCTGCTTTCCCGGACGGCTGTGCCCCATCCG<br>CTTGCTATGTCGACGAGATTCAAGCGTGGTCCACCAATGAACTGACT<br>GTCAATTGGAACTCGGCCCTGTCCTGGGTTGCAAGCTGGGTTGCCG<br>ATCAGGGGAGCGCTGAGCCTGTGCCC |
| Cex<br><br>(**Cex_V1 – 1032nt, 340aa**)<br><br>(Cex_V2 – 1314nt, 438aa) | *cellulomonas fimi*; exo-β-1,4-glucanase, Uniprot: P07986 | Macleod et al. (1994) | ATGCCACGTACCACACCAGCACCAGGTCATCCAGCGCGTGGAGCA<br>CGTACCGCTCTTCGTACGACGCGTCGCCGCGCAGCCACACTGGTC<br>GTGGGTGCCACCGTTGTTTTGCCAGCCCAGGCT**GCCACGACCCTG**<br>**AAAGAAGCCGCCGACGGTGCAGGTCGTGACTTCGGGTTTGCACT**<br>**GGACCCGAATCGTCTTTCGGAGGCACAGTATAAGGCCATTGCAGA**<br>**CTCTGAGTTCAACTTGGTCGTAGCAGAAAACGCAATGAAGTGGGAT**<br>**GCCACGGAACCTAGCCAGAATAGCTTTTCTTTCGGTGCGGGCGAC**<br>**CGTGTGGCGAGCTACGCCGCTGACACCGGAAAAGAACTGTACGG**<br>**CCATACGCTTGTATGGCACTCACAGTTGCCTGACTGGGCTAAGAAT**<br>**CTGAATGGTTCCGCGTTCGAATCAGCAATGGTCAATCATGTCACTA**<br>**AAGTCGCCGACCATTTCGAAGGTAAAGTGGCAAGCTGGGACGTAG**<br>**TAAATGAAGCCTTCGCAGACGGTGACGGACCGCCCCAGGACTCA**<br>**GCTTTTCAGCAAAAATTAGGCAACGGCTATATCGAAACGGCTTTCC**<br>**GTGCTGCTCGTGCGGCAGATCCAACCGCTAAGTTATGTATTAACGA**<br>**CTATAACGTAGAAGGCATCAACGCAAAGTCTAATAGCTTATACGATT**<br>**TAGTCAAGGATTTTAAAGCCCGTGGGGTCCCATTAGATTGTGTAGGT**<br>**TTCCAATCACATCTTATTGTCGGTCAAGTTCCGGGCGATTTCCGCC**<br>**AGAACTTGCAGCGTTTCGCGGACTTGGGGGTCGATGTCCGTATTA**<br>**CGGAGTTGGATATTCGCATGCGTACGCCCTCCGACGCGACAAAG**<br>**CTGGCAACCCAGGCGGCGGATTATAAAAAGGTAGTGCAAGCATG**<br>**CATGCAGGTAACTCGCTGCCAAGGGGTCACGGTCTGGGGCATTA**<br>**CAGATAAGTACAGTTGGGTACCCGACGTTTTTCCCGGAGAAGGAG**<br>**CGGCGCTTGTGTGGGATGCATCTTATGCAAAGAAGCCAGCATACG**<br>**CAGCCGTGATGGAAGCATTCGGCGCGTCCCCTACTCCAACGCCA**<br>**ACGACTCCTACGCCTACCCCAACTACCCCGACCCCAACTCCGAC**<br>**AAGCGGTCCGGCCGGTTGCCAGGTTCTT**TGGGGTGTAAATCAGTG<br>GAATACCGGTTTCACGGCGAACGTTACCGTCAAGAATACCTCTTCAG<br>CTCCTGTAGATGGGTGGACGCTTACGTTCTCTTTCCCTTCGGGCCAG<br>CAAGTCACGCAGGCGTGGTCTTCTACAGTCACACAGAGCGGCTCCG<br>CCGTTACGGTGCGCAATGCCCCTTGGAATGGCTCCATCCCCGCAG<br>GGGGCACCGCCCAATTTGGGTTTAACGGGTCACATACGGGCACCA<br>ACGCTGCCCCGACCGCTTTTAGCTTGAACGGCACGCCATGTACAGT<br>TGGC |

| | | | |
|---|---|---|---|
| E1<br><br>(**E1_V1 – 1074nt, 358aa**)<br><br>(E1_V2 – 1143nt, 381aa) | *Acidothermus cellulolyticus,* endo-β -1,4-glucanase, Uniprot: P54583 | Baker et al. (1994) | ATGCCTCGCGCTCTGCGTCGTGTACCAGGCAGCAGAGTAATGTTGC<br>GCGTTGGTGTGGTAGTTGCTGTTTTAGCGTTGGTTGCGGCTTTGGCCA<br>ATCTGGCAGTCCCACGCCCTGCTCGGGCC**GCAGGCGGGGGTTAT**<br>**TGGCATACGAGTGGCCGTGAAATACTGGACGCAAATAATGTGCCC**<br>**GTCCGTATCGCTGGGATCAACTGGTTTGGGTTCGAAACGTGCAACT**<br>**ACGTTGTTCATGGCTTGTGGAGCCGCGATTATCGCAGCATGTTAGA**<br>**TCAGATCAAGTCCTTGGGATATAATACTATACGTCTTCCCTACTCCG**<br>**ATGATATTCTGAAACCCGGGACGATGCCAAATTCTATCAACTTTTAC**<br>**CAGATGAACCAAGACCTGCAAGGGTTGACCTCCTTGCAAGTCATG**<br>**GATAAGATCGTAGCATACGCGGGCCAGATCGGACTTCGTATAATCT**<br>**TAGATCGCCATAGACCCGATTGTAGCGGTCAGTCTGCCTTGTGGTA**<br>**CACAAGTTCAGTATCAGAAGCAACCTGGATATCTGATTTACAAGCG**<br>**CTTGCACAGCGGTATAAGGGAAACCCTACCGTCGTGGGATTCGAC**<br>**TTACACAATGAGCCTCATGACCCGGCATGTTGGGGCTGTGGTGAC**<br>**CCGTCCATTGATTGGCGTTTGGCAGCGGAAAGAGCTGGGAACGC**<br>**TGTCTTATCAGTGAATCCCAACTTGCTTATCTTCGTTGAAGGCGTTC**<br>**AGTCATACAACGGGGACAGCTATTGGTGGGGCGGCAACCTGCAA**<br>**GGCGCTGGCCAGTACCCCGTAGTACTTAACGTGCCGAACAGACT**<br>**GGTTTATTCCGCCCATGATTATGCAACGAGCGTCTATCCACAAACG**<br>**TGGTTTAGTGACCCTACGTTCCCGAACAATATGCCAGGTATTTGGA**<br>**ACAAAAATTGGGGCTATCTTTTTAATCAGAATATAGCGCCGGTTTGG**<br>**TTAGGTGAGTTTGGGACAACGCTTCAGTCAACCACCGATCAAACTT**<br>**GGCTGAAAACATTAGTACAATATTTGCGGCCGACAGCGCAGTATG**<br>**GTGCAGACAGTTTCCAGTGGACTTTCTGGAGTTGGAATCCGGACTC**<br>**AGGGGATACTGGTGGGATCTTGAAGGACGATTGGCAAACGGTAGA**<br>**TACCGTGAAAGATGGTTACCTTGCTCCGATAAAATCATCGATATTTG**<br>**ACCCGGTT**GGTGCAAGCGCAAGCCCTTCCTCTCAACCCTCACCAA<br>GTGTATCGCCGTCACCGTCTCCCTCCCCCTCAGCCTCTCGCACACC<br>GACGCCGACTCCCACTCCCACTGCGAGCCCCACACCCACATTGAC<br>TCCCACAGCGACCCCAACTCCCACCTAG |
| Endo5A<br><br>(Endo5A – 1053nt, 352aa) | *Paenibacillus sp ICGEB2008*; endo-beta-1,4,-glucanase, Uniprot: G0YA73 | Adlakha et al. (2011) | ATGAAGAAGAAGGGCCTGAAGAAGACCTTCTTTGTCATCGCGAGCCT<br>CGTCATGGGGCTGACCCTCTACGGCTATACCCCGCCGTCCGCGGA<br>CGCCGCCAGCGTCAAGGGGTATTACCACACCCAGGGCAACAAGAT<br>CGTCGATGAGAGCGGCAAGGAGGCGGCCTTCAATGGGCTCAATTG<br>GTTCGGCCTGGAAACCCCCAATTATACCCTGCATGGGCTGTGGTCG<br>CGGTCGATGGATGACATGCTCGACCAAGTCAAGAAGGAGGGGTACA<br>ATCTCATCCGGCTGCCCTACAGCAATCAGCTCTTCGATTCGAGCTCG<br>CGCGCGGATTCCATCGATTACTATAAGAACCCGGATCTGGTCGGCCT<br>GACGCCGATCCAGATCATGGACAAGCTGATCGAGAAGGCCGGCCA<br>GCGTGGCATCCAAATCATCCTGGATCGCCATCGCCCCGGGTCCGG<br>GGGCCAGTCGGAGCTGTGGTATACCTCGCAGTACCCGGAATCCCG<br>GTGGATCTCCGATTGGAAGATGCTCGCCGAACGTTACAAGAATAACC<br>CGACCGTGATCGGGGCGGACCTGCACAACGAGCCCCACGGCCAA<br>GCCAGCTGGGGGACCGGGGATGTGAGCACCGACTGGCGTCTGGC<br>CGCGCAACGTGCGGGCAACGCGATCCTCTCCGTGAACCCGAATTG<br>GCTGATCCTGGTGGAAGGCGTCGACCACAATGTCCAGGGCAACAAT<br>TCCCAGTATTGGTGGGGCGGCAACCTGACCGGCGTGGCCAACTAC<br>CCCGTCGTCCTGGATGTCCCCAACCGGGTCGTCTATTCCCCGCATG<br>ATTACGGGCCGGGGGTGTCGAGCCAACCCTGGTTCAACGACTCGA<br>CGTTTCCGAGCAATCTCCCCGCCATCTGGGACCAAACCTGGGGCTA<br>TATCTCCAAGCAAAACATCGCGCCGGTGCTCGTGGGCGAGTTTGGC<br>GGGCGCAACGTGGATAGCTCGAGCCCGGAAGGCAAGTGGCAAAAT |

| | | | |
|---|---|---|---|
| | | | GCGCTCGTGGACTACATCGGGGCGAACAACCTCTATTTTACCTACTGGTCCCTCAACCCCAATTCGGGCGATACCGGCGGCCTGCTCCTCGACGATTGGGTGACGTGGAATCGCCCGAAGCAAGACATGCTGAGCCGGATCATGAAGCCGGTGAACTTTATCGCGGAACAAGCGAAGGCCTCCGCGGAGTAG |
| Gluc1C<br><br>(Gluc1C – 1335nt, 445aa) | *Paenibacillus sp MTCC 5639*; 1,4-β-glucosidase, Uniprot: P22505 | Adlakha et al. (2012) | ATGAGCGAGAACACGTTCATCTTTCCGGCGACCTTCATGTGGGGGACGTCGACGTCCTCGTACCAGATCGAGGGCGGGACGGACGAAGGCGGGCGCACCCCCAGCATCTGGGATACGTTTTGTCAAATCCCCGGCAAGGTCATCGGGGGCGATTGCGGCGATGTCGCCTGCGATCACTTCCACCACTTTAAGGAGGACGTGCAGCTGATGAAGCAGCTCGGCTTTCTGCATTACCGTTTCAGCGTGGCCTGGCCGCGCATCATGCCGGCCCCGGGCATCATCAACGAGGAAGGCCTCCTGTTCTATGAGCATCTGCTCGACGAAATCGAACTCGCCGGGCTCATCCCCATGCTCACCCTCTATCACTGGGATCTCCCGCAGTGGATCGAGGATGAGGGGGGGTGGACGCAGCGTGAGACGATCCAGCACTTCAAGACCTACGCGAGCGTGATCATGGACCGTTTCGGGGAACGCATCAACTGGTGGAACACGATCAACGAACCGTATTGCGCCTCCATCCTCGGGTATGGGACGGGGGAACATGCCCCCGGCCACGAGAACTGGCGGGAGGCCTTCACCGCCGCCCATCACATCCTGATGTGTCACGGCATCGCCTCGAACCTGCATAAGGAAAAGGGGCTGACCGGGAAGATCGGGATCACCCTCAATATGGAGCACGTGGACGCCGCCAGCGAGCGCCCCGAAGACGTCGCGGCGGCGATCCGCCGTGATGGCTTCATCAACCGTTGGTTTGCGGAACCCCTCTTTAACGGGAAGTACCCGGAAGATATGGTCGAGTGGTACGGCACCTACCTCAACGGGCTGGACTTTGTGCAGCCCGGCGATATGGAACTGATCCAACAACCCGGCGATTTTCTGGGCATCAATTACTACACGCGTAGCATCATCCGGTCGACCAATGACGCGTCCCTCCTCCAGGTGGAGCAAGTCCACATGGAGGAACCCGTCACCGACATGGGCTGGGAGATCCATCCCGAGTCGTTCTATAAGCTCCTCACCCGCATCGAGAAGGACTTCAGCAAGGGCCTCCCCATCCTCATCACCGAAAACGGGGCCGCGATGCGCGATGAGGTGGTGAACGGCCAGATCGAAGACACCGGCCGCCAGCGTTATATCGAAGAGCATCTGAAGGCGTGTCACCGTTTCATCGAAGAGGGGGGGCAACTCAAGGGGTATTTCGTCTGGTCGTTTCTCGACAACTTTGAGTGGGCGTGGGGGTACAGCAAGCGCTTCGGGATCGTGCATATCAATTATGAAACGCAAGAGCGGACCCCGAAGCAATCCGCCCTCTGGTTTAAGCAAATGATGGCCAAGAACGGGTTCTAA |

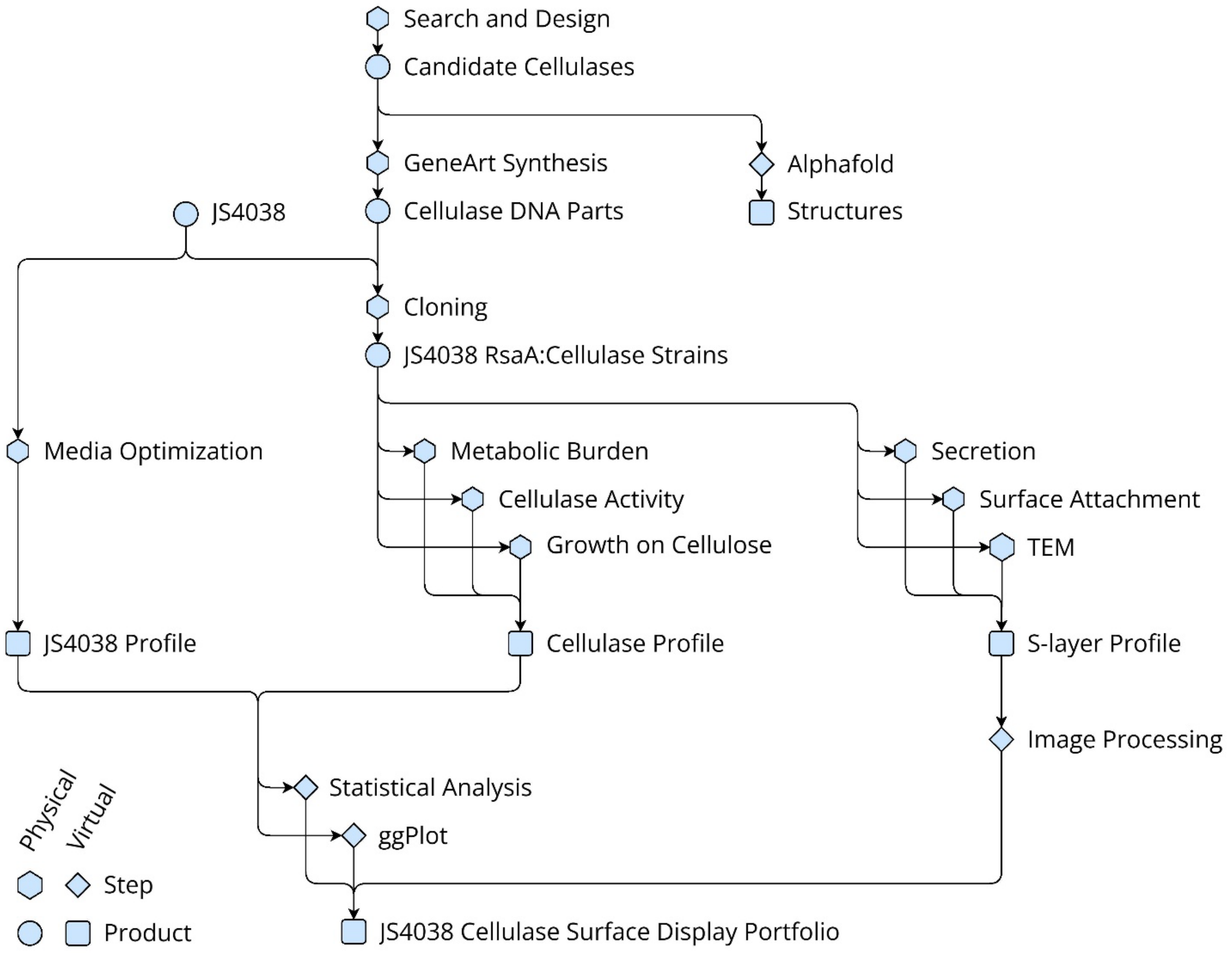


**Figure S1.** Workflow of methods employed in this study, including physical and virtual steps and products. Made using Drawio (www.drawio.com).

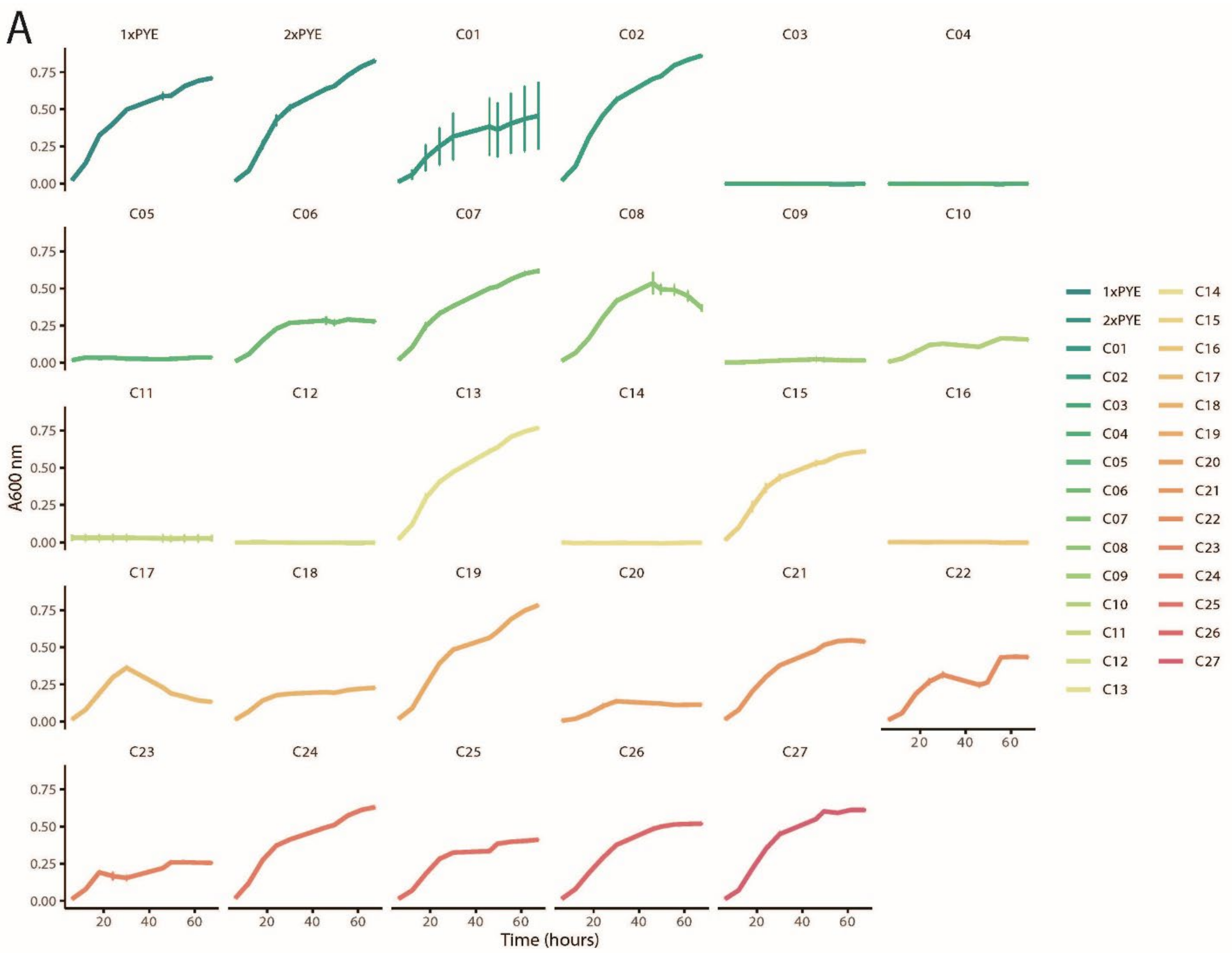


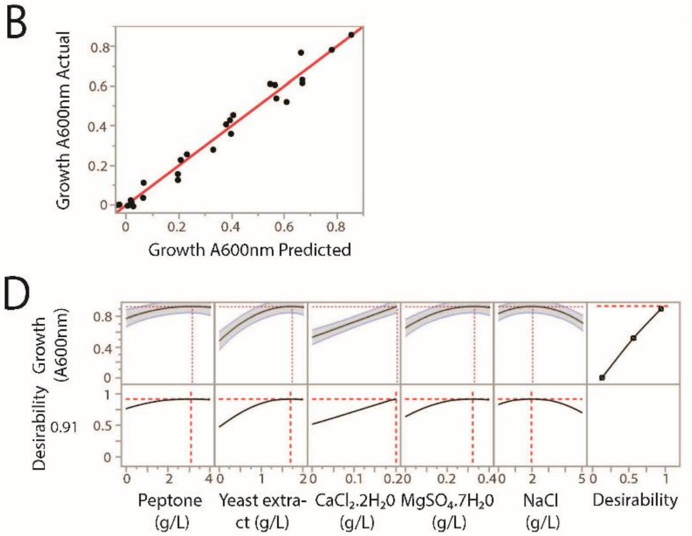


C

| Term | Estimate | Std Error | Prob>\|t\| |
|---|---|---|---|
| Intercept | 0.6830195 | 0.034923 | <.0001* |
| Yeast extract(0,2) | 0.1480656 | 0.016934 | <.0001* |
| CaCl2.2H20(0,0.2) | 0.0990733 | 0.017179 | <.0001* |
| MgSo4.7H20(0,0.4) | 0.0968309 | 0.016804 | <.0001* |
| Yeast extract* CaCl2.2H20 | 0.0849098 | 0.019052 | 0.0006* |
| Peptone(0,4) | 0.0563339 | 0.017306 | 0.0063* |
| Peptone* CaCl2.2H20 | 0.0472638 | 0.019041 | 0.0275* |
| CaCl2.2H20* MgSo4.7H20 | 0.0350763 | 0.019041 | 0.0884 |
| CaCl2.2H20* NaCl | 0.0279771 | 0.018881 | 0.1622 |
| Peptone* Yeast extract | -0.045522 | 0.019206 | 0.0339* |
| Yeast extract* NaCl | -0.047488 | 0.018969 | 0.0264* |
| NaCl(0,5) | -0.057667 | 0.0164 | 0.0038* |
| Peptone* Peptone | -0.065128 | 0.035799 | 0.092 |
| MgSo4.7H20* MgSo4.7H20 | -0.107825 | 0.039293 | 0.0167* |
| NaCl* NaCl | -0.149532 | 0.037427 | 0.0015* |
| Yeast extract* Yeast extract | -0.155057 | 0.038744 | 0.0015* |

**Figure S2**. Surface Response Model of JS4038 growth for PYE medium optimisation. (A) Growth curves measured at 600 nm absorbance ($A_{600}$) for JS4038 across 27 media compositions generated using a five-factor central composite design in JMP, alongside standard 1× PYE and 2× PYE controls. Factor concentrations ranged from 0× to 2× levels reported in published PYE formulations. Curves represent mean $A_{600}$ absorbance from triplicate 200 µL cultures, with error bars representing standard deviation. (B) A scatterplot showing the relationship between experimentally measured $A_{600}$ values (actual) and the model-predicted $A_{600}$ values derived from the quadratic response-surface model. Each black point represents an individual experimental condition. The red diagonal line indicates perfect agreement between predicted and observed values ($R^2$ = 0.96). (C) Final parameter estimates from the response surface model. The table lists coefficients, standard errors, and significance values for all retained terms. Positive coefficients represent factors and interactions that increase predicted $A_{600}$, while negative coefficients indicate inhibitory effects. Coefficients are ordered by estimate from positive to negative. (D) Prediction profiler showing cross-sections of the multivariate response surface. Desirability functions (0–1) can be applied to maximise $A_{600}$, identifying optimal medium concentrations at desirability = 1.

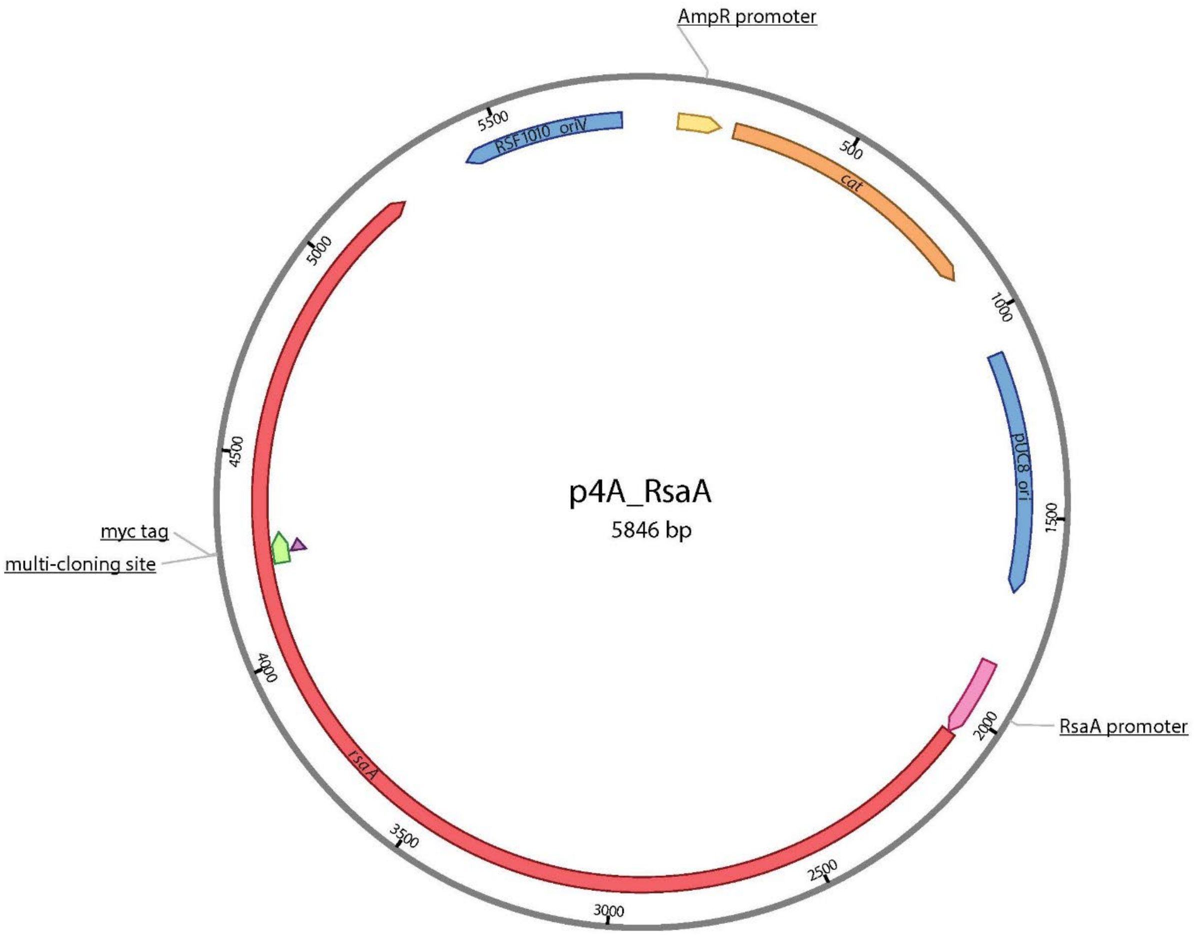


**Figure S3**. Plasmid map of p4A_RsaA723 with selected feature information. p4A_RsaA723 has two origins of replication including pUC8 for maintenance in *E. coli*, and oriV for maintenance in *Caulobacter vibrioides*, in addition to a chloramphenicol acetyltransferase (*cat*) antibiotic selection marker, and a multi-cloning site (MCS) embedded within the S-layer gene *rsaA* supporting in-frame insertions.

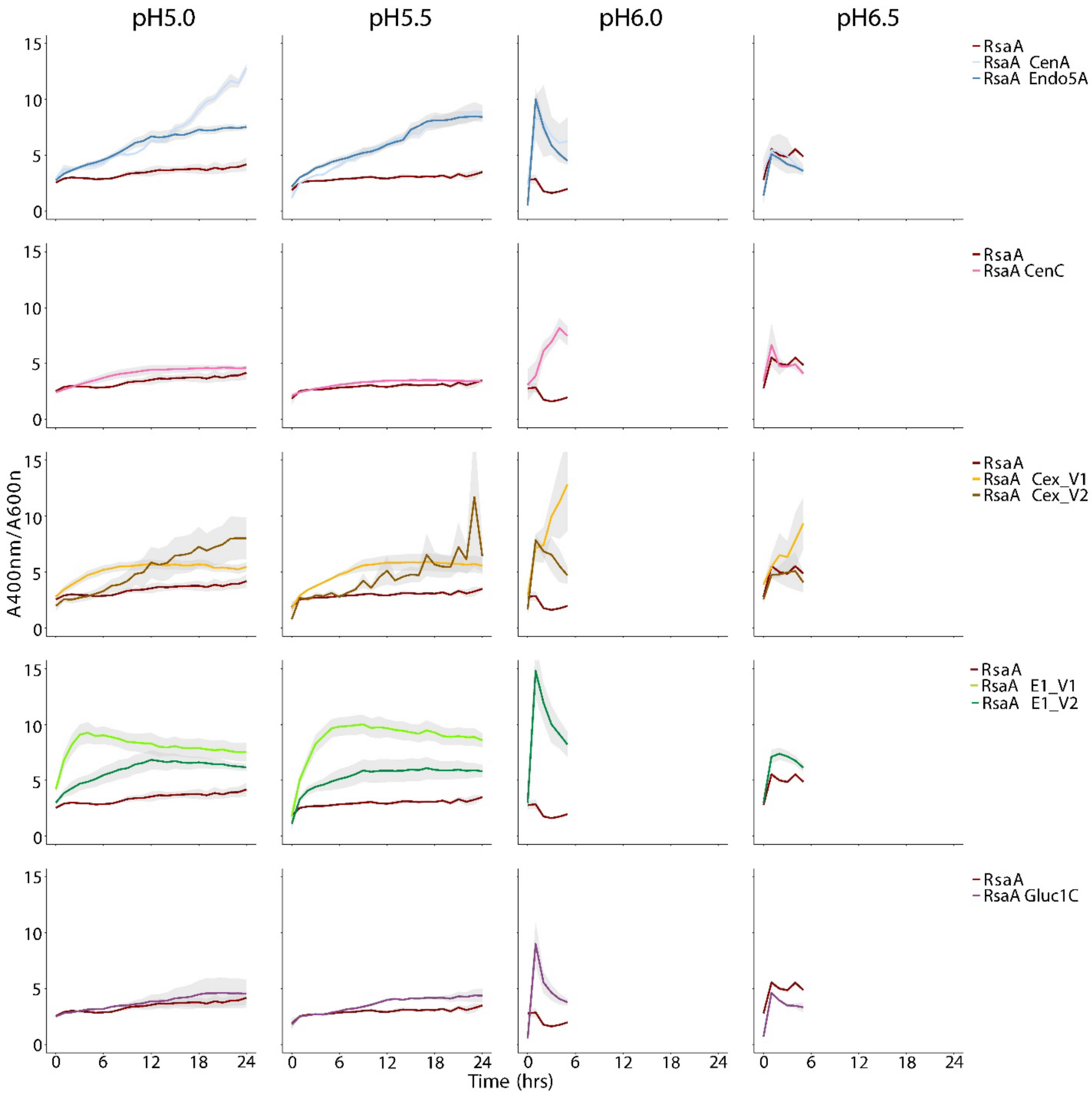


**Figure S4**. All DNP-C experiments performed during this study at four different pH levels. Experiments were performed for 24hrs at pH5 and pH5.5, and 6hrs at pH6 and pH6.5. Absorbance at 400 nm ($A_{400}$) was corrected for cell density using $A_{400}$. Shaded error bars represent standard error of the mean of three biological replicates.

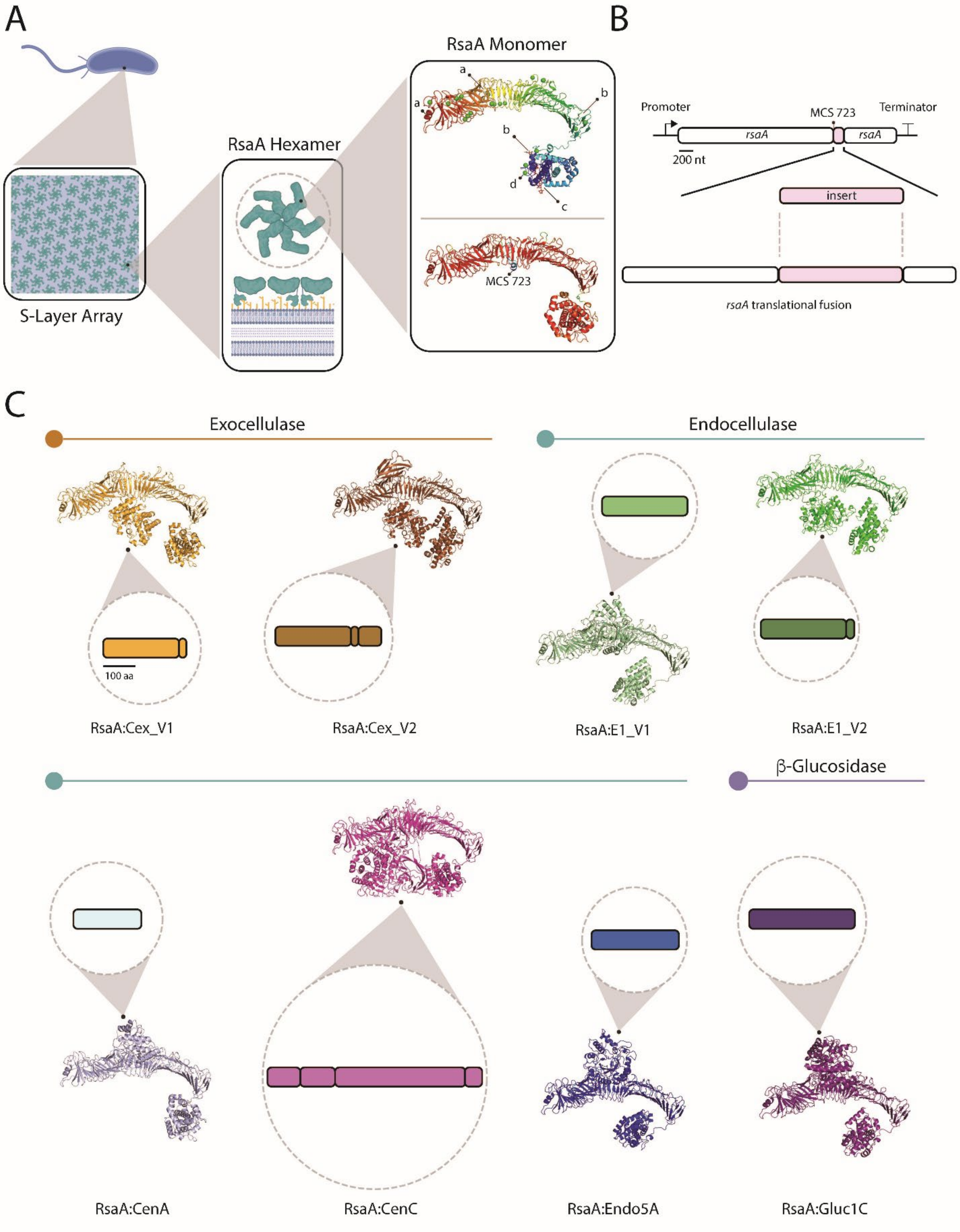

A
RsaA Monomer
RsaA Hexamer
S-Layer Array
a
a
b
b
d
c
MCS 723
B
Promoter
MCS 723
Terminator
rsaA
rsaA
200 nt
insert
rsaA translational fusion
C
Exocellulase
Endocellulase
100 aa
RsaA:Cex_V1
RsaA:Cex_V2
RsaA:E1_V1
RsaA:E1_V2
β-Glucosidase
RsaA:CenA
RsaA:CenC
RsaA:Endo5A
RsaA:Gluc1C

**Figure S5**. Concept map and modeled proteins employed in this study. (A) Schematic of the S-layer array, its hexamer unit, and two structural models of an RsaA monomer. The top model depicts experimentally resolved RsaA (PDB: 6Z7P); Inter-unit contact sites for hexamer to hexamer-lattice assembly (a), intra-unit contact sites for monomer to hexamer assembly (b), the N-terminal domain's LPS binding site to enable surface attachment of the S-layer (c), and the $Ca^{2+}$ ions it complexes (d), are labeled. The bottom model depicts an AlphaFold predicted structure of RsaA with the multi-cloning site at amino acid 723 (labeled). (B) Conceptual framing of the cloning workflow during this study to produce S-layer fusion proteins. (C) Schematics of each cellulase part cloned into the S-layer gene, with associated Alphafold predicted structural models.

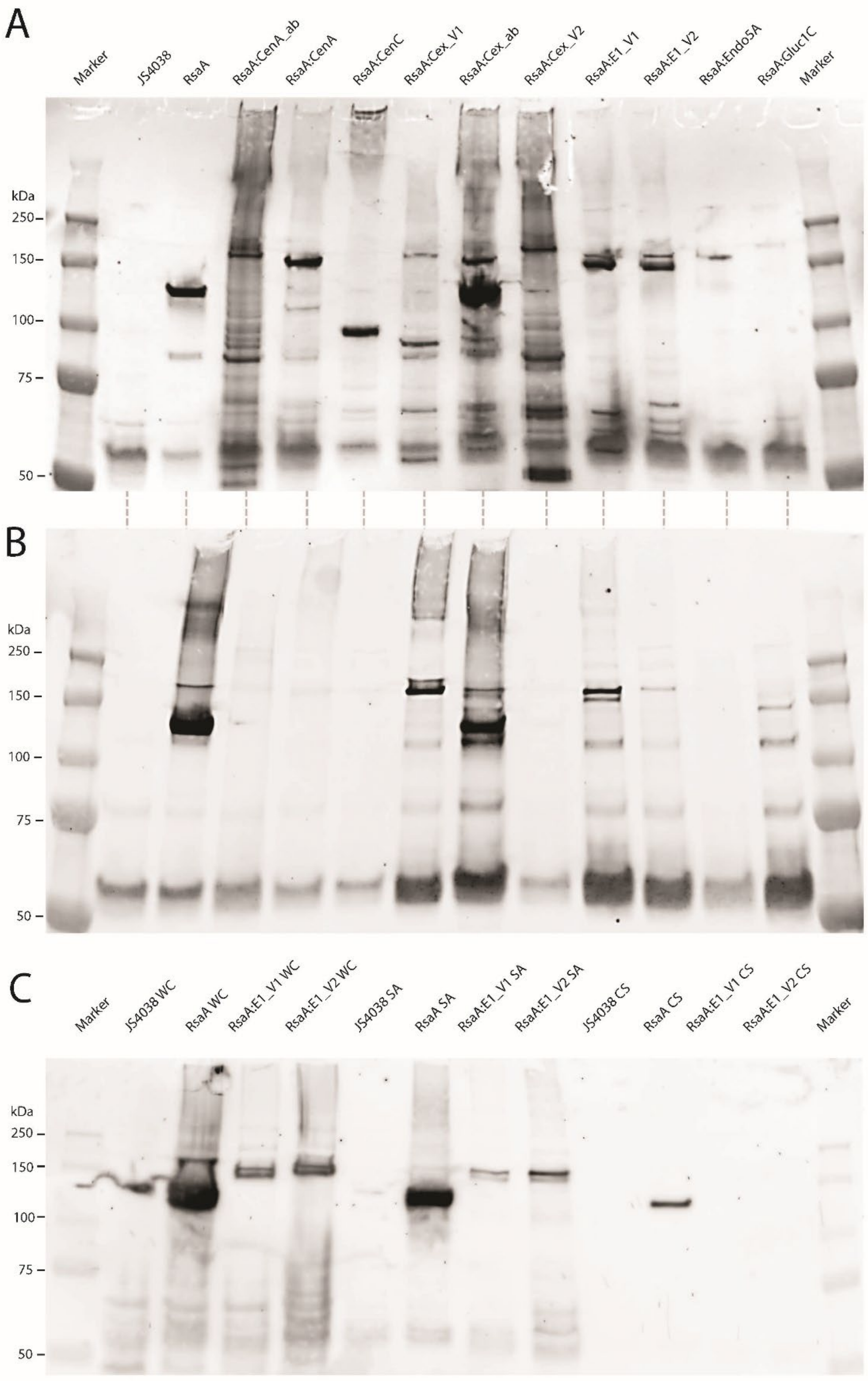

A
Marker
JS4038
RsaA
RsaA:CenA_ab
RsaA:CenA
RsaA:CenC
RsaA:Cex_V1
RsaA:Cex_ab
RsaA:Cex_V2
RsaA:E1_V1
RsaA:E1_V2
RsaA:Endo5A
RsaA:Gluc1C
Marker
kDa
250
150
100
75
50
B
kDa
250
150
100
75
50
C
Marker
JS4038 WC
RsaA WC
RsaA:E1_V1 WC
RsaA:E1_V2 WC
JS4038 SA
RsaA SA
RsaA:E1_V1 SA
RsaA:E1_V2 SA
JS4038 CS
RsaA CS
RsaA:E1_V1 CS
RsaA:E1_V2 CS
Marker
kDa
250
150
100
75
50

**Figure S6.** Immunoblots conducted for this study. (A) Immunoblot with polyclonal anti-RsaA antibodies against whole-cells of JS4038 cropped to five markers. RsaA:CenA_ab and RsaA:Cex_ab contain abandoned constructs not used throughout the rest of the study. (B) Immunoblot with polyclonal anti-RsaA antibodies against surface attached protein fractions of JS4038 cropped to five markers. (C) Immunoblot with polyclonal anti-RsaA antibodies against whole-cells of JS4038 (lanes 2-5), surface attached protein fractions of JS4038 (lanes 6-9), and against JS4038 culture supernatant (lanes 10-13) cropped to five markers.

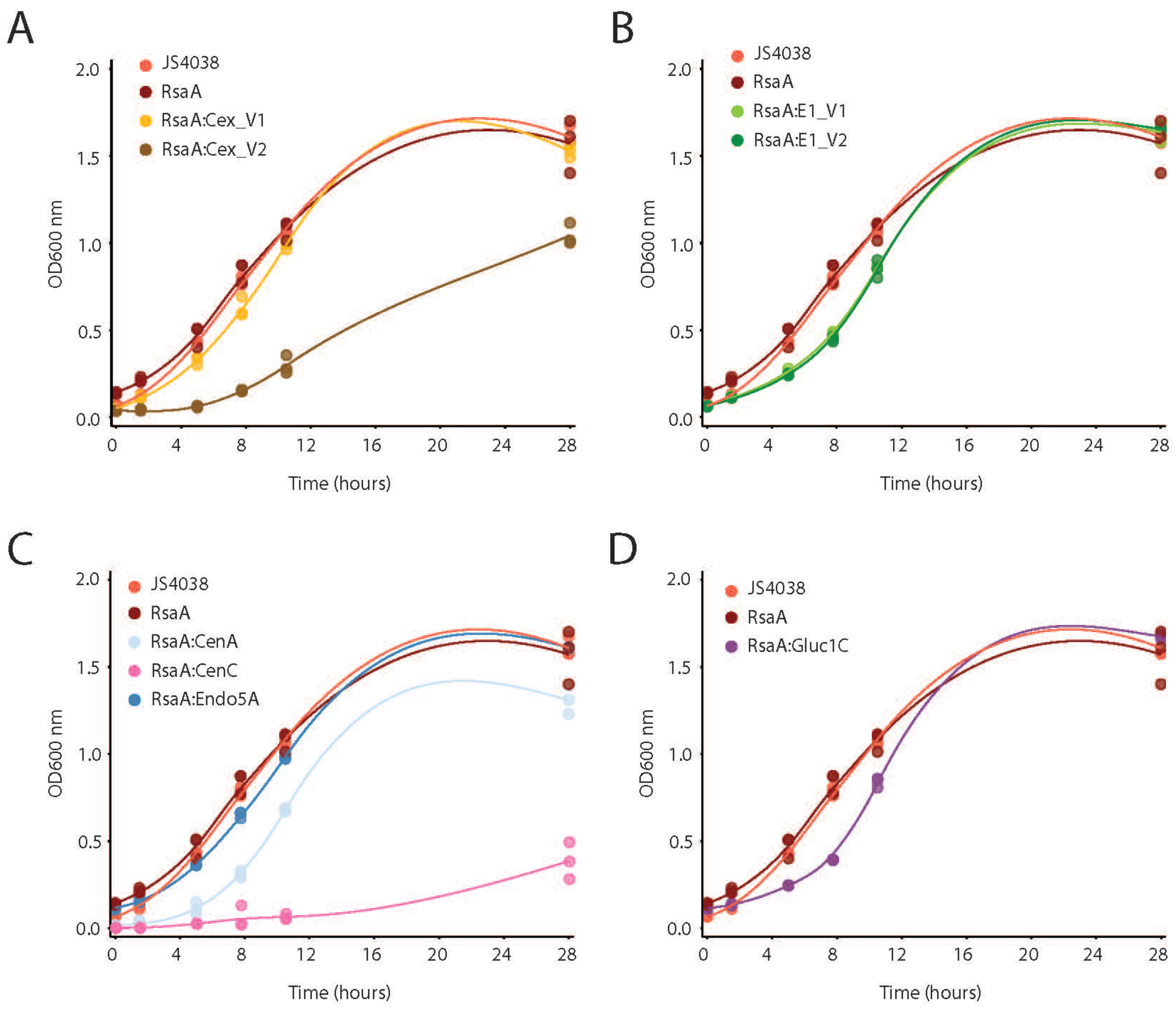


**Figure S7.** (A-D) Growth curves ($OD_{600}$) of JS4038 with and without expressing RsaA or an RsaA:Cellulase fusion protein. Points represent individual replicates; lines represent a LOESS-smoothed best-fit line.

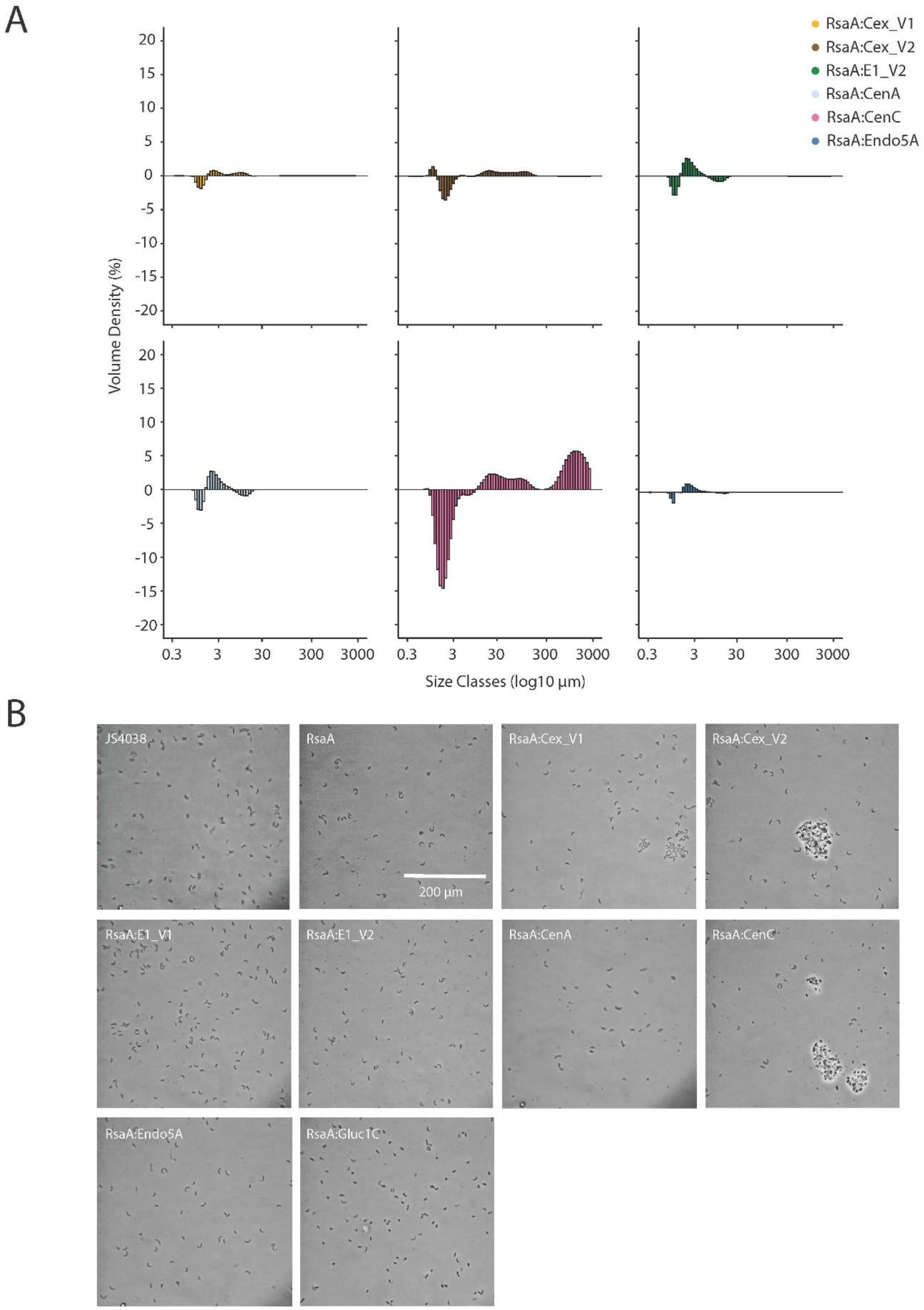
A
RsaA:Cex_V1
RsaA:Cex_V2
RsaA:E1_V2
RsaA:CenA
RsaA:CenC
RsaA:Endo5A
Volume Density (%)
Size Classes (log10 µm)
B
JS4038
RsaA
RsaA:Cex_V1
RsaA:Cex_V2
200 µm
RsaA:E1_V1
RsaA:E1_V2
RsaA:CenA
RsaA:CenC
RsaA:Endo5A
RsaA:Gluc1C

**Figure S8.** Cell clumping phenotype analysis of JS4038 with and without expressing RsaA or an RsaA:Cellulase fusion protein. (A) Average % volume density of four replicates of cell clump size in each culture compared to cells expressing RsaA only (% volume density of fusion strain minus % volume density of positive RsaA control) measured with a MasterSizer instrument. (B) Brightfield phase contrast microscopy images of each culture (43,000x).

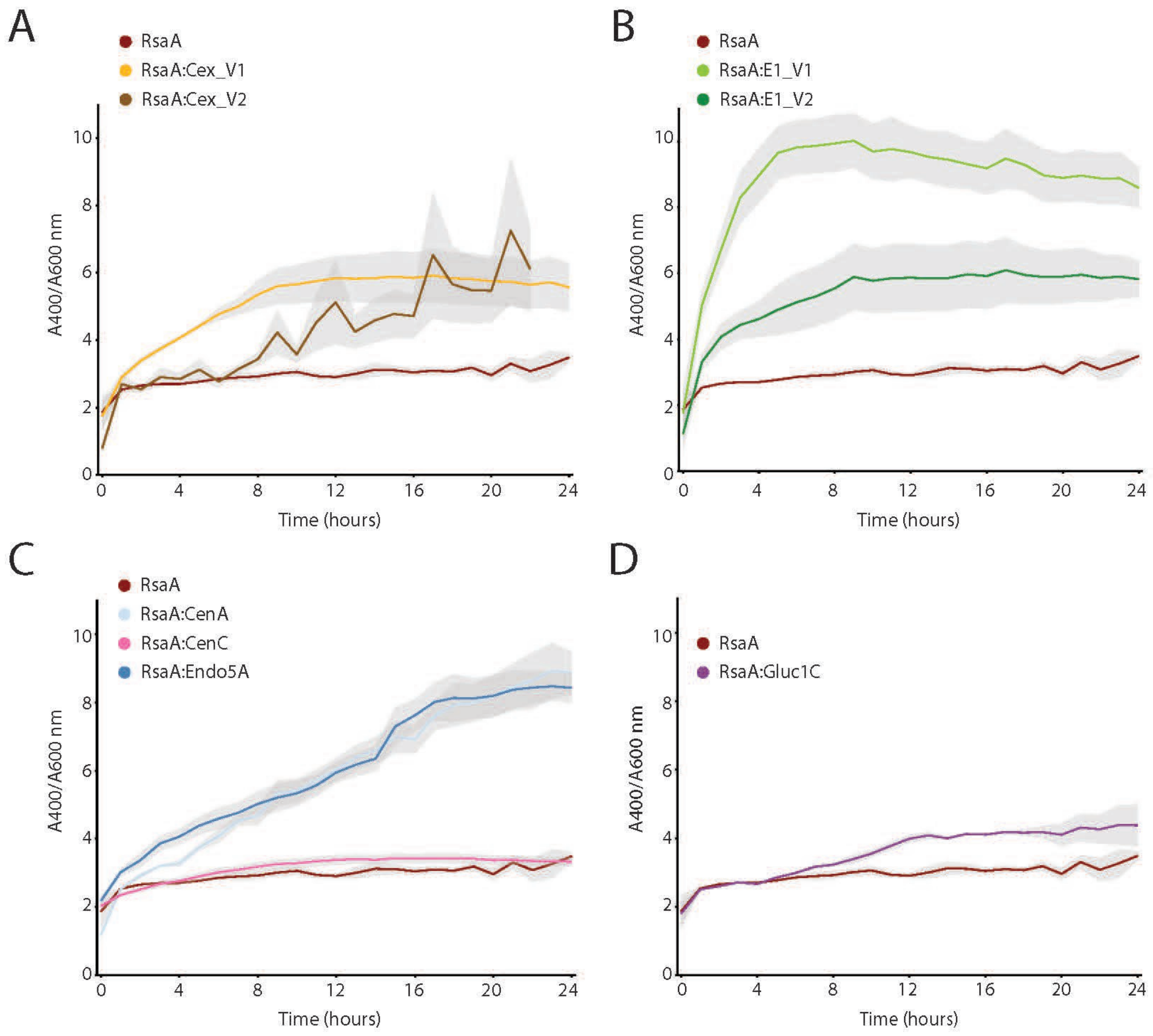


**Figure S9.** (A-D) DNP-C experiments performed during this study at pH5.5 on JS4038 expressing RsaA or an RsaA:Cellulase fusion protein. Absorbance at 400 nm ($A_{400}$) was corrected for cell density using $A_{600}$. Shaded error bars represent standard error of the mean of three biological replicates.

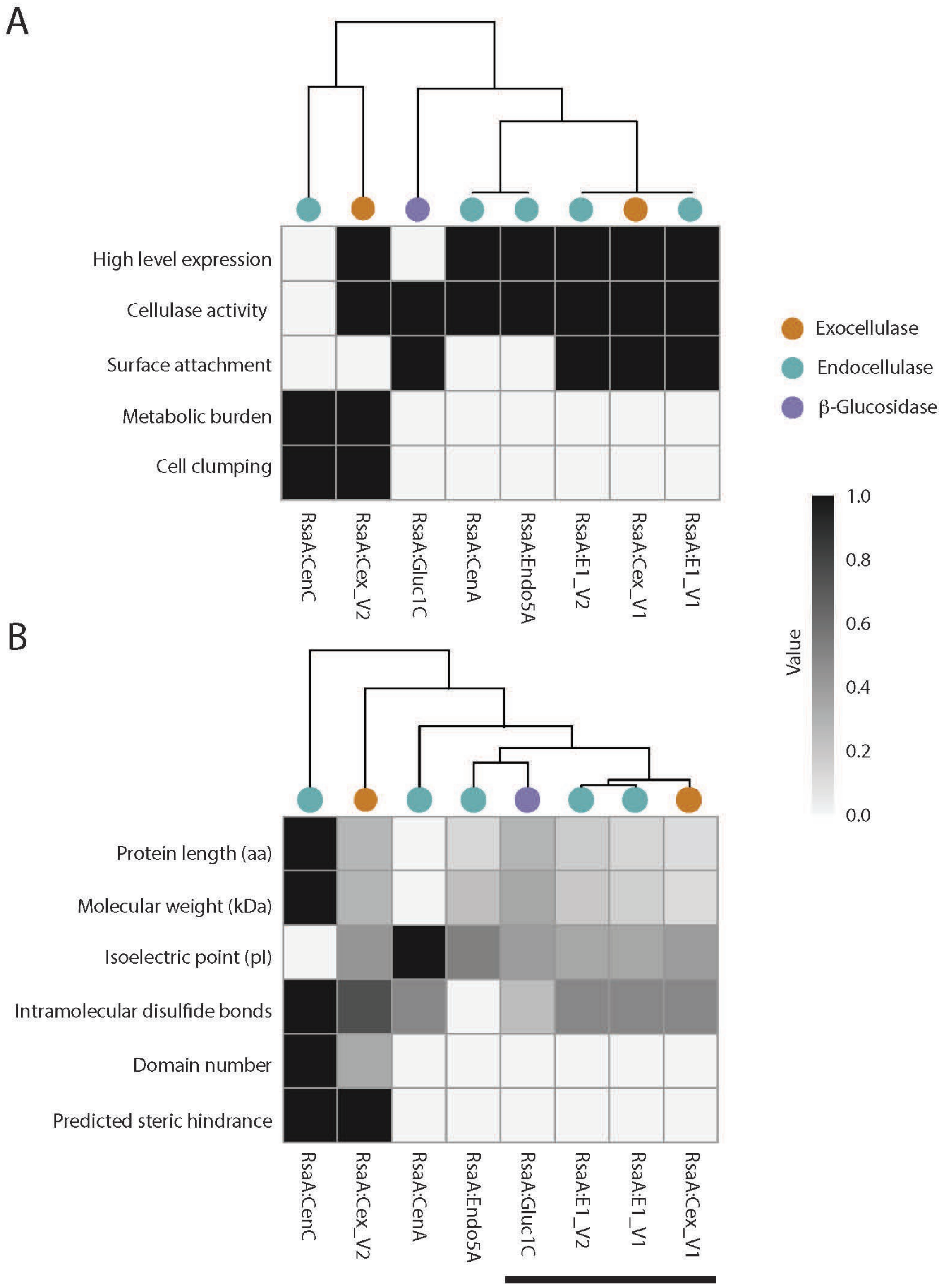
A
High level expression
Cellulase activity
Surface attachment
Metabolic burden
Cell clumping
RsaA:CenC
RsaA:Cex_V2
RsaA:Gluc1C
RsaA:CenA
RsaA:Endo5A
RsaA:E1_V2
RsaA:Cex_V1
RsaA:E1_V1
Exocellulase
Endocellulase
β-Glucosidase
Value
1.0
0.8
0.6
0.4
0.2
0.0
B
Protein length (aa)
Molecular weight (kDa)
Isoelectric point (pI)
Intramolecular disulfide bonds
Domain number
Predicted steric hindrance
RsaA:CenC
RsaA:Cex_V2
RsaA:CenA
RsaA:Endo5A
RsaA:Gluc1C
RsaA:E1_V2
RsaA:E1_V1
RsaA:Cex_V1

**Figure S10.** (A) Hierarchically clustered heatmap of phenotypic and functional traits across RsaA-cellulase fusion constructs displayed in this study. Input data represent binary values (1 = present, black, 0 = absent, white) for five phenotypes across eight constructs. Rows and columns are clustered based on similarity to highlight patterns of co-occurrence and divergence among traits, informing construct performance. (B) Hierarchically clustered heatmap of protein design features across the same fusion constructs. Numeric input values for six features were independently normalized per row from 0 to 1. This row-wise normalization enables comparison of relative feature magnitudes across constructs. Clustering illustrates relationships among constructs based on design attributes.

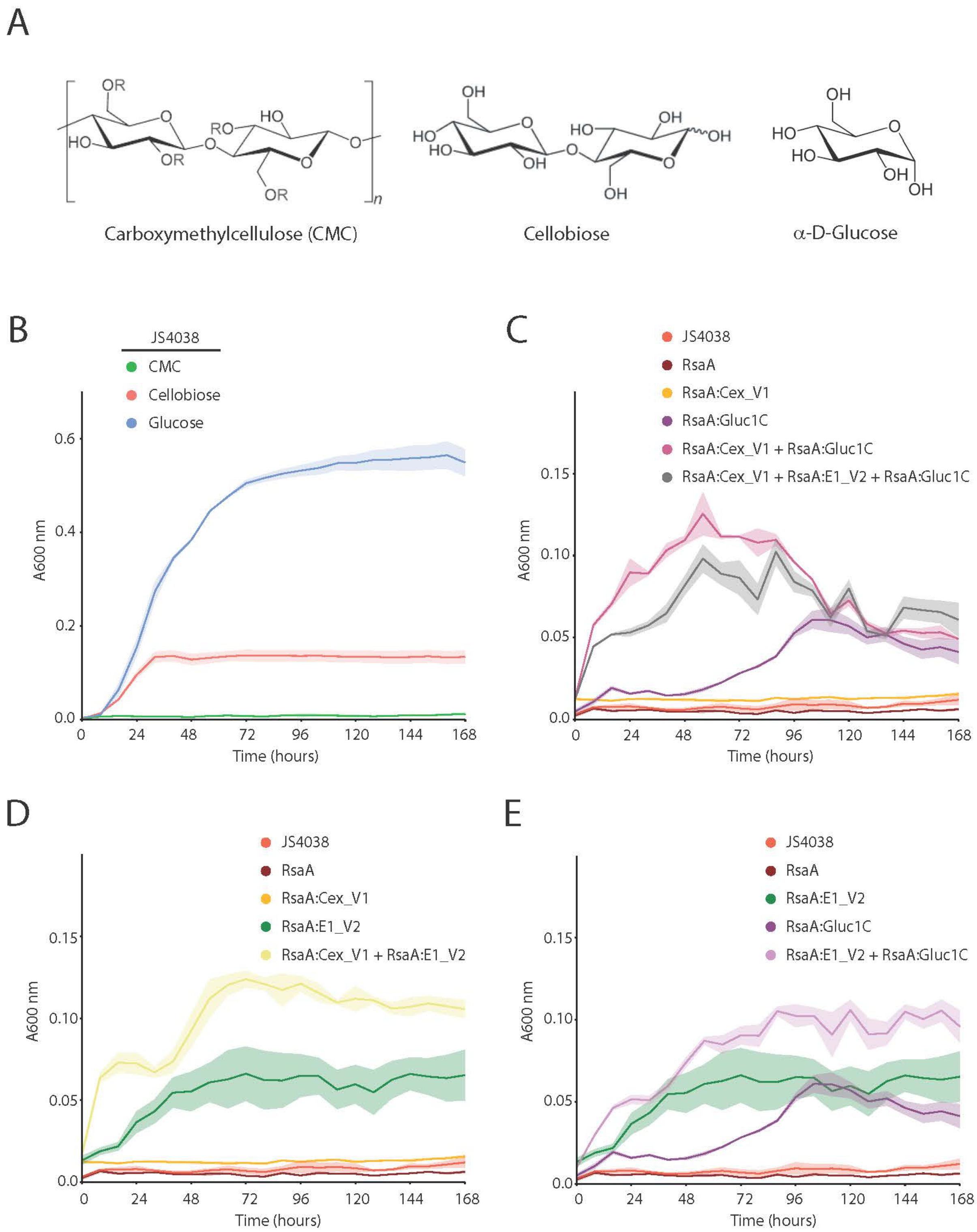
A
Carboxymethylcellulose (CMC)
Cellobiose
α-D-Glucose
B
JS4038
CMC
Cellobiose
Glucose
A600 nm
Time (hours)
C
JS4038
RsaA
RsaA:Cex_V1
RsaA:Gluc1C
RsaA:Cex_V1 + RsaA:Gluc1C
RsaA:Cex_V1 + RsaA:E1_V2 + RsaA:Gluc1C
A600 nm
Time (hours)
D
JS4038
RsaA
RsaA:Cex_V1
RsaA:E1_V2
RsaA:Cex_V1 + RsaA:E1_V2
A600 nm
Time (hours)
E
JS4038
RsaA
RsaA:E1_V2
RsaA:Gluc1C
RsaA:E1_V2 + RsaA:Gluc1C
A600 nm
Time (hours)

**Figure S11.** (A) Molecular diagrams of the three carbon sources utilised in the cellulose growth experiment. (B) Growth curves ($A_{600}$) of JS4038 in HMG minimal media with glucose, cellobiose or carboxymethyl-cellulose (CMC) as the sole carbon source. (C-E) Growth curves ($A_{600}$) of JS4038 with and without expressing RsaA or an RsaA:Cellulase fusion protein in HMG minimal media with CMC as the sole carbon source. Strains were assayed independently, mixed with one other strain 50:50, or two other strains 33:33:33. Shaded areas represent standard error of the mean of three biological replicates.
